# Impacts of reduced inorganic N:P ratio on three distinct plankton communities in the Humboldt upwelling system

**DOI:** 10.1101/550046

**Authors:** Kristian Spilling, Maria-Teresa Camarena-Gómez, Tobias Lipsewers, Alícia Martinez-Varela, Francisco Díaz-Rosas, Eeva Eronen-Rasimus, Nelson Silva, Peter von Dassow, Vivian Montecino

## Abstract

The ratio of inorganic nitrogen to phosphorus (NP) is projected to decrease in the Eastern Boundary Upwelling Systems (EBUS) due to warming of the surface waters. In an enclosure experiment, we employed two levels of NP ratios (10 and 5) for three distinct plankton communities collected along the coast of central Chile (33°S). The primary effect of the NP treatment was related to different concentrations of NO_3_, which directly influenced the biomass of phytoplankton. Additionally, low inorganic NP ratio reduced the seston NP and Chl *a*-C ratios, and there were some effects on the plankton community composition, e.g. benefitting *Synechococcus* spp in some communities. One of the communities was clearly top down controlled and trophic transfer to grazers was up to 5.8% during the 12 day experiment. Overall, the initial plankton community composition was more important for seston stoichiometry and trophic transfer than the inorganic NP ratio. Any long term change in the plankton community structure will likely have greater impact than direct effects of a decreasing inorganic NP ratio on the Humboldt Current ecosystem.

## Introduction

The Humboldt Current, one of the four major Eastern Boundary Upwelling Systems (EBUS), transports water from the Sub-Antarctic zone northwards along the western cost of South-America before it deflects off the coast by Ekman transport. Cold, nutrient-rich sub-surface water replaces the surface water mass in several upwelling centers along the coast of Chile and Perú. The high concentration of inorganic nutrients in the upwelling water supports high primary productivity, which form the basis for one of the richest fisheries in the world (Montecino and Lange 2009).

One of the emerging changes to ocean chemistry is the loss of dissolved oxygen (DO), driven by increasing temperature causing reduced gas solubility and ventilation of the deep ocean (Keeling et al. 2010; Stramma et al. 2010). The DO dynamics is also governed by respiration, and the absence of DO opens up a range of niches for microbial life using alternative electron acceptors, with direct implications for cycling of key elements, e.g. C, N, P and Fe (Kirchman 2012). Expanding Oxygen Minimum Zones (OMZs) in the EBUS regions will consequently change the chemistry of the upwelling water with potential implications for biological production (Keeling et al. 2010; Schmidtko et al. 2017).

Phosphate, the bioavailable form of P, is not a redox-sensitive substance, but there is a strong, negative correlation between DO and phosphate concentrations due to the coupling between P and Fe cycling (Einsele 1938). Under anoxic conditions, phosphorus bound to oxidized metal compounds (e.g. Fe^3+^) is released during reduction processes. Continuing expansion of the South Pacific OMZ towards the shelf will likely increase the release of P into the upwelling water in the Humboldt Current.

The effect of expanding OMZs on N cycling is more complex. The main reactive inorganic nitrogen (N_r_) species: nitrate, nitrite and ammonium, have dual roles; they can be used as nutrients for phytoplankton and bacteria, or in microbial redox-processes that end in the formation of N_2_ gas, which is not readily available for biological uptake (with the exception of N-fixing diazotrophs). Denitrification and anaerobic oxidation of ammonium with nitrite (anammox) are major N_r_ removing processes and the OMZs are major N_r_ sinks globally (Kuypers et al. 2005). However, both these processes are ultimately driven by organic matter oxidation, which could function as a negative feedback mechanism, i.e. if lower N_r_ in the upwelling water reduces the organic N input to the OMZs (Capone and Hutchins 2013; Kalvelage et al. 2013). The opposite process, N-fixation, transforms N_2_ into organic N by marine diazotrophs and provides a source of “new” N_r_ to the ecosystem. N-fixation can take place both within the OMZs and in the surface waters (Fernandez et al. 2011; Hamersley et al. 2011). There is likely not a direct correlation between the OMZ size and N_r_ loss (Yang et al. 2017). However, given the magnitude of N_r_ loss in the OMZs, its projected expansion and model simulation of N sources and sinks, the bioavailable N_r_ in the upwelling water will most likely decrease, meaning a reduction in the supply of N_r_ (Kalvelage et al. 2011; Landolfi et al. 2013). Loss of DO may consequently affect both N and P cycling in opposing direction, decreasing N_r_ while increasing bioavailable P.

Upwelling off the Chilean coast takes place north of 38°S and extends seasonally down to 46°S (Strub et al. 2019). Large chains of diatoms typically dominate during periods with strong upwelling, whereas picophytoplankton, nanoplankton, and dinoflagellates increase in importance during periods with less frequent or lower intensity upwelling (Anabalón et al. 2007; Collado-Fabbri et al. 2011; Anabalón et al. 2016). The bacterial community composition is known to be influenced by the DO concentration (Aldunate et al. 2018), and the plankton community structure have implications for trophic transfer pathways through the food web, and export of organic material (Vargas et al. 2007; Ochoa et al. 2010). Given the importance of plankton community composition, it is essential to understand the consequences of a reducing inorganic NP ratio for different communities as the effect of a decreasing inorganic NP ratio will likely be modified by the phytoplankton community composition.

Although the Humboldt current system has been studied over decades, there are in general few experimental studies from the region, in particular addressing changing nutrient stoichiometry, with a few notable exceptions: Franz et al. (2012) and Hauss et al. (2012) reported from the same shipboard, mesocosm experiment off the coast of Peru using four different inorganic NP ratios ranging from 2.5 to 16. They found a direct correlation between the inorganic NP ratio and particulate organic NP ratio after N depletion but with a lower limit of 5 (PON:POP ratio) (Franz et al. 2012), and the biomass was related to the amount of inorganic N addition. There were some changes to the community structure, e.g. low NP ratio benefitting *Heterosigma* sp. and *Phaeocystis globosa* (Hauss et al. 2012). Hauss et al. (2012) did one additional experiment with and without mesozooplankton and inorganic NP ratios of 20, 3.4 and 2.8, where the inclusion of mesozooplankton from the start reduced the phytoplankton biomass peak due to grazing.

The plankton community composition can be more important than direct changes in environmental conditions, e.g. ocean acidification (Eggers et al. 2014). Any effects of changes to the NP ratio will likely depend on the structure of the plankton community. In order to address the combined effect of community composition and reduction in the inorganic NP ratio, we carried out a short-term experiment using three distinct plankton communities and two different NP ratios. The experiment was set up with water from three different locations off the Chilean coast, and we applied NP ratios 10 and 5. A NP ratio of 10 corresponds closely to the mean inorganic NP ratio integrated over the upper 30 m during upwelling periods in the study area (Anabalón et al. 2016), while a NP ratio of 5 was chosen to correspond to a future scenario with reduced inorganic NP ratio in the upwelling water. Based on the previous studies from the region (Franz et al. 2012; Hauss et al. 2012), our original hypotheses were: (1) there is a clear effect of inorganic NP on seston stoichiometry; and (2) changes in the inorganic NP ratio will shift the community composition. In addition we wanted to compare the variability caused by inorganic NP ratio and community composition and hypothesized that (3) the variability in seston stoichiometry is greater between different communities than between the inorganic NP ratio treatments.

## Methods

### Experimental set-up

Water was collected from three different locations along the Chilean coast referred to as ‘North’, ‘Central’ and ‘South’ (Fig 1) during austral summer (March). The sampling sites were chosen based on different hydrological conditions (Wieters et al. 2003; Narváez et al. 2004), with the aim of having different initial plankton community composition. Communities ‘North’ and ‘Central’ originated from frequent upwelling sites with low temperatures (<13°C), whereas the community ‘South’ was collected in the vicinity of the Maipo River outlet, but with similar salinity. Water was collected from 5 m depth using a Niskin water sampler. Temperature varied between 12.2-13.4 °C and salinity between 34.5-34.6 (Table 1). The water was sieved through a 200 µm mesh to remove mesozooplankton to reduce immediate top down control. The water containers were covered with black plastic in order to prevent excessive light during the transport. The initial sampling (day 0) for plankton community composition (microzooplankton, phytoplankton and bacteria) was taken from the collected water before the addition of inorganic nutrients (day 1). A seawater sub-sample (60 mL) was frozen (−20°C) for later measurement of initial nutrient content.

**Table 1.** The coordinates of the sampling locations (Fig 1), the physical characteristics of the water and the NP ratio after nutrient additions.

|  | Location | Coordinates | Salinity | Temperature | Initial nutrients |  | NP ratio after nutrient addition |  |
| --- | --- | --- | --- | --- | --- | --- | --- | --- |
|  |  |  |  |  | NO<sub>3</sub> | PO<sub>4</sub> | NP10 | NP5 |
| 862 | North | 33.18611S-71.71933W | 34.6 | 12.3 °C | 4.7 µM | 1.0 µM | 10.3 | 3.3 |
| 863 | Central | 33.35088S-71.68208W | 34.6 | 12.2 °C | 24.8 µM | 2.6 µM | 11.4 | 6.7 |
| 864 | South | 33.61597S-71.64649W | 34.5 | 13.4 °C | 7.3 µM | 1.4 µM | 9.6 | 3.6 |

**Fig 1.**
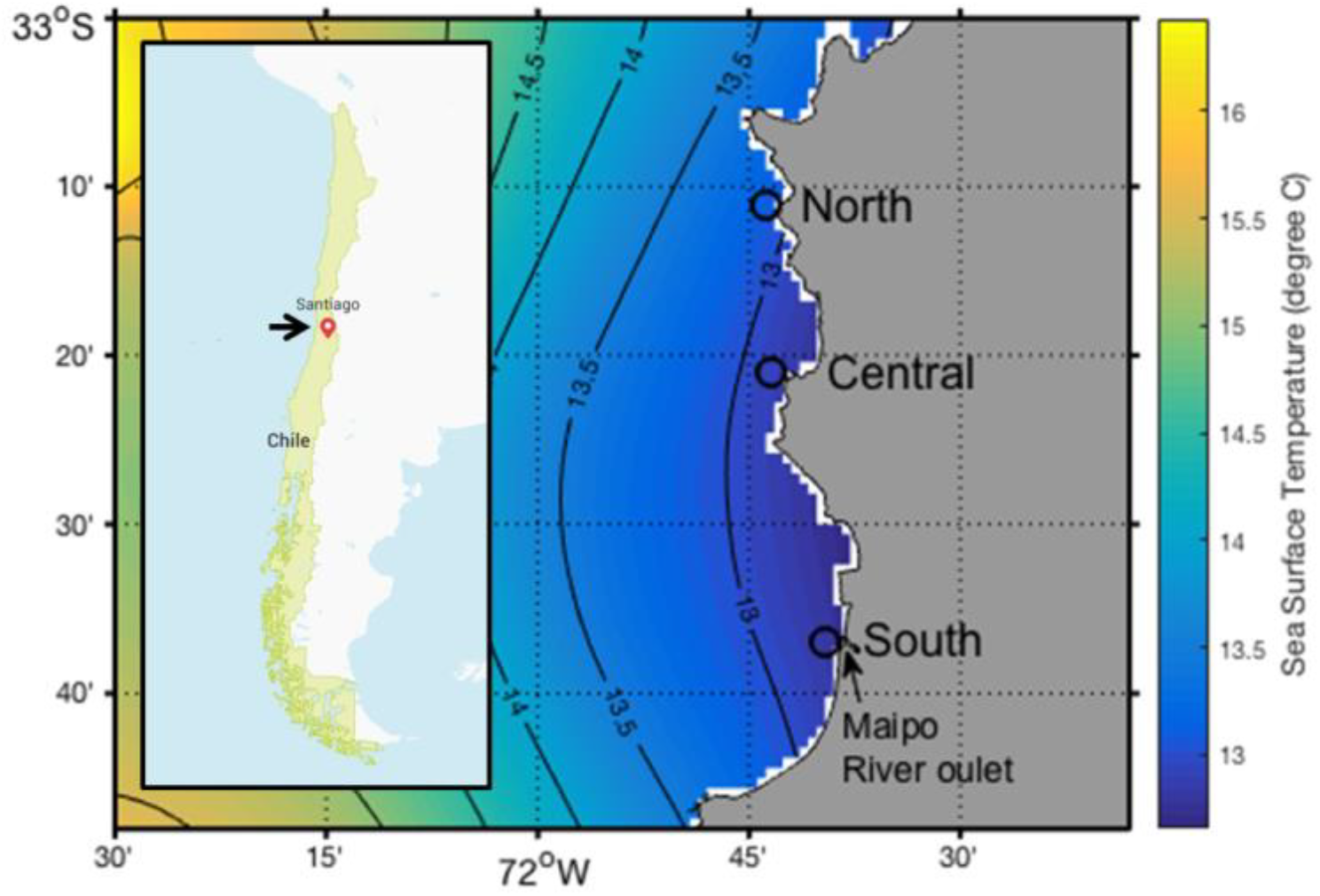
Map with the composite sea surface temperature (°C) at the day of water collection (13 March 2014) calculated from satellite observations (Multi-scale ultra-high resolution sea surface temperature, NASA https://mur.jpl.nasa.gov/). The arrow in the inserted map of Chile indicate the Maipo River outlet. The water was collected from three different sites: ‘North’, ‘Central’ and ‘South’ (open circles).

The sampled water was set up in clean, 15 L carboys at the Las Cruces Marine Biological Station. To clean the carboys, they were first filled with artificial salt water (deionized water with added salt, 35g L^-1^) for a week at room temperature. After that the carboys were emptied, acid washed (4 h in 10% HCl), rinsed 6 times with deionized water and dried. Before the beginning of the experiment, the carboys were filled with the water collected from the three different locations. The water was collected from 5 m depth with a Niskin bottle, and the three replicate carboys were filled consecutively. Before the start of the experiment the carboys were submerged in a water bath placed outside under natural sunlight. The water bath was lined with a black plastic in order to limit backscattering of light. There was continuous flow through of sea water, pumped from 5 m depth outside the station, in the water bath keeping the temperature relative stable at 14-16 °C. A neutral screen covered the water bath reducing the ambient irradiance, and the light measured with a spherical light collector (Waltz) inside the carboys was 10% of the light above the shading screen. There were continuous measurements of light and temperature (every 30 s), which is presented in the supplementary material (supplementary figures S1 and S2). Light was measured with a LiCor cosine collector connected to a data logger (MadgeTech). Temperature was measured and stored using an Ebro EBI310 thermometer with integrated data logger, connected to a dipping probe placed in the water bath. Two thermometers were used, measuring the temperature both at the inflow and outflow end of the water bath.

The experiment was set up in a 3^2^ design (3 communities, 2 NP levels) with 3 replicates of each combination, totaling 18 carboys. In order to avoid any confounding effect of the placement of the carboys in the water bath, caused by slight variation in temperature and light intensity, we used a combination of random and consecutive placement. In short, all of the first replicates were randomly distributed, followed by the second and third replicates also with random distribution within each group. The rationale behind this was to avoid the possibility of having all of the three replicates placed close to each other in the water bath, which is a possibility when using full randomization of placement. There was continuous, gentle bubbling with pre-filtered (0.2 µm) air to provide gas exchange and preventing settling. The outflowing air was also going through a 0.2 µm filter to prevent contamination.

We were not able to determine the inorganic nutrient concentration before the start of the experiment, and could not adjust the inorganic nutrient addition according to the ambient concentrations. Inorganic N and P were added to a final concentration of 14.4 and 3.6 µmol NO_3_ L^-1^ and 0.9 and 1.6 µmol PO_4_ L^-1^ for the high and low NP ratio respectively (Table 1). For simplicity we refer to the NP treatments as NP10 and NP5 throughout the text, which was the average high and low NP ratio. In order to prevent any other nutrient limitation (e.g. silicate limitation of diatoms or iron limitation), dissolved silicate, trace metals and vitamins were added to all treatments using F/2 medium stock (Guillard 1975) to a final concentration in all treatments fixed to the N addition in the NP10 treatment (e.g. 7 µmol L^-1^ DSi was added to all treatments). After the experiment, the initial nutrient concentrations (day 0) were determined (Table 1), together with sub-samples taken during the experiment (day 1 to day 12).

### Sampling

The experimental containers were placed inside the water bath on 13 March 2014 (day 0) and sub-samples were taken for filtration and determining the initial plankton communities. Inorganic nutrient were added before sampling in the morning (08:00) of day 1, and sampling continued until day 12. Sub-samples (5 ml) were taken daily for measurement of fluorescence properties. Other measurements were done every two or three days (500 ml) with the exception of phytoplankton and bacteria community samples, which were collected three times during the experiment (additional 500 ml).

Fluorescence was measured after dark acclimation for 15 min, in a 10 mm quarts cuvette with an AquaPen (Photon Systems Instruments) fluorometer using 450 nm excitation light. The minimum fluorescence (F_0_) was used as a proxy for Chl *a* concentration. The maximum (F_m_) and variable (F_m_-F_0_ = F_v_) and fluorescence was used to calculate the photochemical efficiency (F_v_/F_m_).

### Inorganic nutrients and organic elements

For determination of inorganic nutrients (NO_3_, PO_4_ and DSi), samples were stored in pretreated bottles, cleaned with 5% HCl and dried with acetone, and placed in a freezer (−20°C). The nutrient concentrations of all samples were determined right after the experiment. Samples were thawed quickly at 40°C and nutrients were determined with an autoanalyzer according to Atlas et al. (1971), at the Escuela de Ciencias del Mar, Pontificia Universidad Católica de Valparaíso.

Sub-samples were filtered onto GF/F filters to determine Chl *a* and particulate organic nutrients. For Chl *a*, the filters were subsequently flash frozen in liquid nitrogen and kept frozen until Chl *a* was extracted by adding 10 mL ethanol. The vials were left in room temperature for 24 h before the Chl *a* concentration was determined using a fluorescence spectrophotometer (Cary Eclipse, Agilent Technologies) calibrated against known Chl *a* standards (Sigma-Aldrich). For particular organic carbon (POC), nitrogen (PON) and phosphorus (POP), acid washed and pre-combusted (450°C for 4 h) GF/F filters were used. Filters were allowed to dry and stored at room temperature until determination of particular nutrients. POC and PON were determined by mass spectrometer (Europa Scientific, ANCA-MS 20-20) and POP by the methods described in Solórzano and Sharp (1980). Due to a failure of the mass spectrometer, some samples were lost for day 9 data, preventing calculation of error estimates for POC and PON for some treatments at that time point.

There was no direct measurement of dissolved organic nitrogen (DON) or phosphorus (DOP). The relative change was, however, calculated according to:

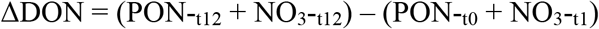

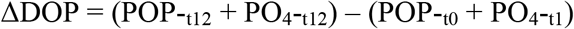

The particulate organic fraction was taken from day 0 and day 12 (t0 and t12) and inorganic nutrients after their addition on day 1 (t1) and from the last day (t12).

### Phytoplankton community composition

The phytoplankton community was determined by flow cytometry (BD InFlux Cell Sorter equipped with a 488 nm and 640 nm lasers), FlowCam (FluidImaging) and light microscopy. The flow cytometry samples were taken at all sampling time points, and 1.2 ml were preserved with 120µl Glutaraldehyde (25%, quality grade), mixed and flash-frozen with liquid nitrogen before storage at −80°C until cells were enumerated. Chlorophyll-containing cells were identified by red fluorescence (692 nm/40 nm bandpass) and phycoerythrin-containing cells were identified by orange fluorescence (580 nm/30 nm bandpass). Nanophytoplankton were distinguished from picoplankton based on light scatter (FSC and SSC) and pulse width (transit time) compared to 3 µm UltraRainbow fluorescent particles (Spherotech).

The FlowCam samples were taken three times: before the beginning of the experiment (day 0), at the end (day 12) of the experiment and at the Chl *a* maximum based on fluorescence measurements (days 5, 7 and 9 for communities ‘North’, ‘Central’ and ‘South’ respectively). For these samples, 50 mL were preserved with acid Lugol’s solution (0.5% v/v) and stored at 4°C until they were counted. The samples were left for a few hours at room temperature before a minimum of 1000 cells (>2 µm) were counted with the FlowCam. The biovolume was determined automatically by the software from the pixels of individual images of different cells and then sorted into groups by a combination of the sorting options provided by the software and manual classification. In addition all samples from day 12 were counted using an inverted microscope (Letiz Labovert). A minimum of 400 cells were counted using 400x magnification.

### Bacterial community composition

The samples for bacterial community composition were collected before the beginning of the experiment (day0), close to the Chl *a* peak (day 5, 7 and 9 for communities ‘North’, ‘Central’ and ‘South’ respectively), and in the end of the experiment (day12). The samples (500 ml) were filtered onto 0.2 µm sterile Whatman cellulose ester filters (Whatman) and stored at −80 °C. The DNA extraction was done using a Power Soil DNA isolation kit (Mo Bio Laboratories Inc.). The 16S ribosomal RNA (rRNA) gene V1 to V3 hypervariable region was amplified in two polymerase chain reactions (PCR), using the universal bacterial primers F8 and R492. Illumina MiSeq paired-end multiplex sequencing was performed at the Institute of Biotechnology, University of Helsinki, Finland. Primer removal was done with Cutadapt (v. 1.10 with Python 2.7.3, (Martin 2011)).

In total, ∼6.2 million raw reads of the 16S rRNA gene were obtained. The pair-end reads were merged using PEAR software (v 0.9.6, (Zhang et al. 2013)). The UPARSE pipeline(Edgar 2013) was used for quality filtering (> 400 base pairs (bp), maximum expected error 1), chimera checking(Edgar et al. 2011) and operational taxonomic unit (OTU) clustering (97%, (Edgar 2013)). In total, 1.9 million merged sequences were obtained after the quality filtering. Taxonomic classification of the OTUs was done with Silva (v. 119, 60 % confident threshold(Quast et al. 2012) in Mothur 1.38.1.1, (Schloss et al. 2009)). After the removal of chloroplasts, mitochondria and singletons, 927 OTUs with 1 million sequences were obtained for further analyses.

### Zooplankton enumeration

Microzooplankton were enumerated by the FlowCam from the same samples as the phytoplankton community described above. Mesozooplankton that had developed during the experiment (initially they were removed), was enumerated on the last day. Four liters of sample water was sieved through a 200 µm net, and placed on a GF/F filter. Copepods were enumerated under a stereo microscope (Leitz) and the POC and PON from the >200 µm fraction, were subsequently determined from the filters as described above. We did not observe any large diatom chains (or other autotrophs) on the filters.

The trophic transfer of N (TR_N_) into mesozooplankton was calculated according to:

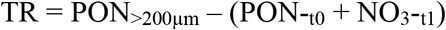

where PON_>200µm_ is the PON concentration in the >200µm fraction on the last day, the PON_-t0_ is the total PON concentration at the start of the experiment and NO_3-t1_ the concentration of NO_3_ after nutrient addition.

### Data treatment and statistics

Treatment effects were statistically analyzed with two-way analysis of variance (ANOVA) using Tukey’s post-hoc test. The two tested variables were location and NP treatment, and the test was done by pairwise comparison. Fluorescence, Chl *a* and POC were transformed to a cumulative value, adding each value to the previously determined value. A linear regression was done for all replicates, and the slope of the regression was used for the two-way ANOVA. This way the temporal aspect was included into the analysis. For the phytoplankton and bacteria communities, we did a repeated measures permutational ANOVA (PERMANOVA) with pairwise comparisons (Anderson 2001), on the data determined on the last day of the experiment. The data were standardized using logarithmic transformation (Anderson et al. 2006) and square-root transformation on the raw data for the phytoplankton community and bacterial community, respectively. The factors analyzed, treatment (NP5: n = 18 and NP10: n = 18) and communities (North, Central, South) were set as fixed-factors. A total of 9999 permutations, using unrestricted permutation of raw data (Manly 2006), were performed, which is recommended when sample size is small (Anderson et al. 2008). The homogeneity of dispersion (i.e. homogeneity of variance) was tested with permutational multivariate analysis of dispersion (PERMDISP (Anderson 2006)), using the distance to the centroids. To test the treatment effect on specific communities, a one way ANOVA was applied.

The phytoplankton and bacterial community dynamics were constructed and visualized using non-metric multidimensional scaling (NMDS) plots. All multivariate analyses and NMDS plots were performed on the Bray-Curtis dissimilarity matrix. The figures were done in R (R CoreTeam, 2016) using the R packages vegan v. 2.3-2. For the multivariate analyses we used PRIMER v. 6 software with the add-on package permutational ANOVA/MANOVA+ (PERMANOVA+).

## Results

### Initial conditions

At the time of water collection for setting up the experiment, the Chl *a* concentration of the initial phytoplankton communities: ‘North’, ‘Central’ and ‘South’ varied (Fig. 1.), but none of these phytoplankton communities were nutrient depleted. The community ‘North’ had the highest Chl *a* concentration (27.4 µg Chl *a* L^-1^) and lowest nitrate concentration (4.7 µmol NO_3_ L^-1^), the community ‘Central’ had the lowest Chl *a* (3.5 µg Chl *a* L^-1^) and highest nitrate concentration (24.8 µmol NO_3_ L^-1^) and the community ‘South’ had intermediate Chl *a* (15.3 µg Chl *a* L^-1^) and nitrate concentration (7.3 µmol NO_3_ L^-1^). After the carboys had been filled, inorganic N (NO_3_) and P (PO_4_) were added, producing an average NP ratio of 10 (‘NP10’) and 5 (‘NP5’; Table 1).

### Development of inorganic nutrients

Inorganic nitrate was rapidly taken up after the beginning of the experiment (Fig 2), and was depleted already at day 3 for treatments: ‘North-NP10’, ‘North-NP5’ and ‘South-NP5’. For treatments ‘South-NP10’ and ‘Central-NP5’, nitrate was depleted at day 5, and for ‘Central-NP10’ at day 7.

**Fig 2.**
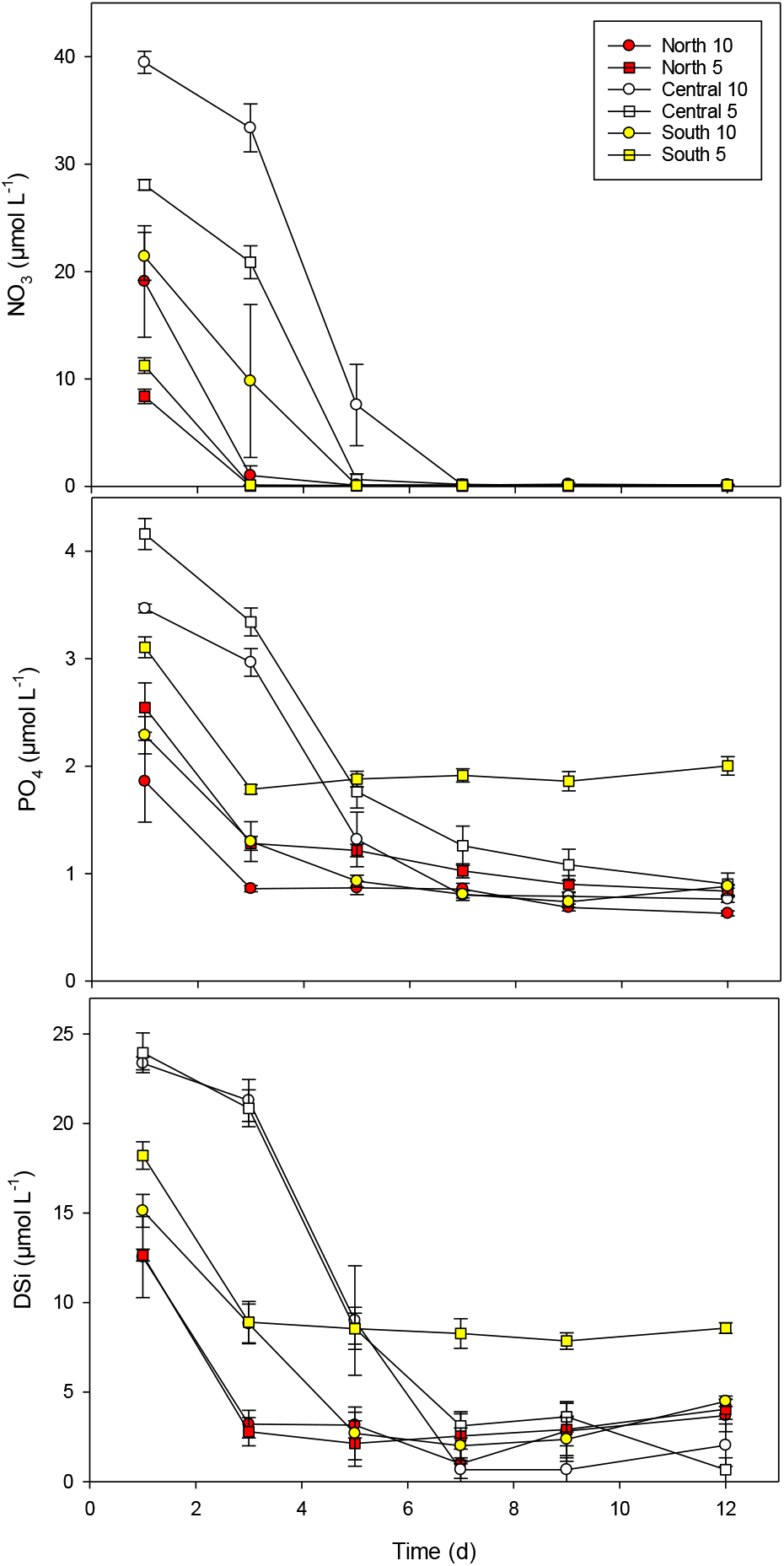
Concentration of inorganic nutrients throughout the experiment. ‘North’, ‘Central’ and ‘South’ represents the three communities (Fig 1), with 2 levels of added NP ratios (‘NP10’ and ‘NP5’, Table 1). Error bars are SE (n = 3).

Phosphate decreased rapidly from the initial values of 2-4 µmol PO_4_ L^-1^, but was not depleted in any of the treatments (Fig. 2). There was approximately 1 µmol PO_4_ L^-1^ at the final day of the experiment (day 12) in all treatments except for ‘South-NP5’, where the average phosphate concentration was ∼2 µmol PO_4_ L^-1^ (Fig 2). For communities ‘North’ and ‘Central’, the difference in PO_4_ L^-1^ between ‘NP10’ and ‘NP5’ decreased from 0.7 µmol PO_4_ L^-1^ initially to 0.2 and 0.1 µmol PO_4_ L^-1^ respectively at day 12. For the community ‘South’, the difference in PO_4_ concentration between ‘NP10’ and ‘NP5’ prevailed throughout the experiment.

The community ‘Central’ had the highest initial concentration of Dissolved Silicate (DSi; 24 µmol L^-1^), but it decreased to the lowest concentration of the three communities during the experiment (1-2 µmol DSi L^-1^). The community ‘South’ had intermediate DSi concentration initially (17 µmol DSi L^-1^) and treatments ‘South-NP10’, ‘North-NP10’ and ‘North-NP5’ were similar in terms of the concentration after nitrate depletion (∼4 µmol DSi L^-1^). The treatment ‘South-NP5’, had the highest residual DSi concentration at the end of the experiment (∼9 µmol DSi L^-1^).

### Organic nutrient pools

The particulate organic nutrient pools, together with the inorganic nutrients, suggested clear differences in the dissolved organic nitrogen (DON) and phosphorus (DOP) pools when comparing the drawdown of NO_3_ and PO_4_ with the increase in particulate organic nitrogen (PON) and phosphorus (POP) respectively (Tables 2 and 3). The calculated DON was different among the communities (two-way ANOVA, *p* <0.001), but there was no effect of the inorganic NP ratio (two-way ANOVA, *p* = 0.85). Using this approach of closing the N and P budget, the communities ‘North’ and ‘South’ consumed 13.3 ± 2.0 (SE) and 2.5 ± 2.9 (SE) µmol DON L^-1^, respectively, whereas the community ‘Central’ produced 6.9 ± 1.1 (SE) µmol DON L^-1^ (Table 2). For DOP there was no treatment effect (two-way ANOVA, p = 0.45 and *p* = 0.93 for the community and the NP treatment, respectively) and there was an average increase in the DOP pool of 0.83 ± 0.11 (SE) µmol DOP L^-1^ over the course of the experiment (Table 3).

**Table 2.**
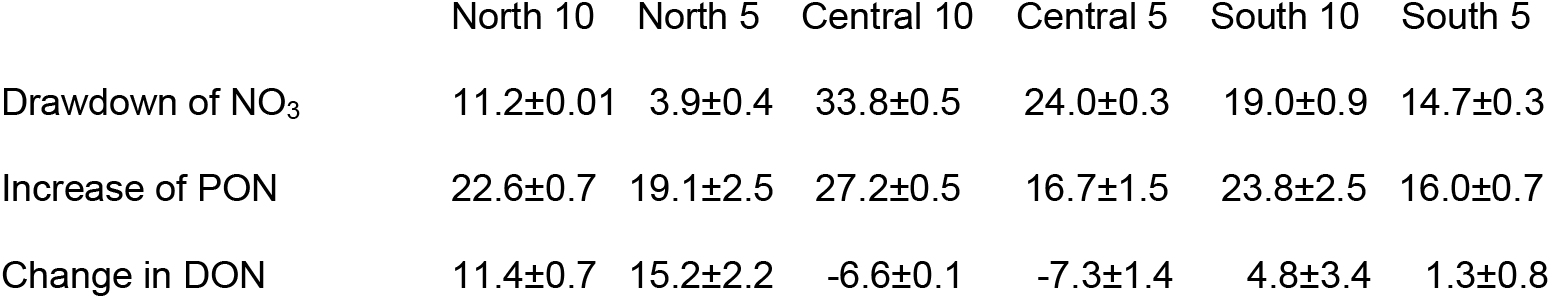
The dissolved organic nitrogen (DON) was calculated by comparing the drawdown of NO_3_ (concentration at T0 minus concentration at T12) with the increase in particulate organic nitrogen (PON) (concentration at PON maximum minus concentration at T0). The calculated change in DON is the difference between increase in PON and drawdown of NO_3_, either suggesting uptake or production (negative values) of DON. All values in μmol L-1, error estimates are given as ± Standard Error.

| 875 |  | North 10 | North 5 | Central 10 | Central 5 | South 10 | South 5 |
| --- | --- | --- | --- | --- | --- | --- | --- |
| 876 | Drawdown of NO<sub>3</sub> | 11.2±0.01 | 3.9±0.4 | 33.8±0.5 | 24.0±0.3 | 19.0±0.9 | 14.7±0.3 |
| 877 | Increase of PON | 22.6±0.7 | 19.1±2.5 | 27.2±0.5 | 16.7±1.5 | 23.8±2.5 | 16.0±0.7 |
| 878 | Change in DON | 11.4±0.7 | 15.2±2.2 | -6.6±0.1 | -7.3±1.4 | 4.8±3.4 | 1.3±0.8 |

**Table 3.**
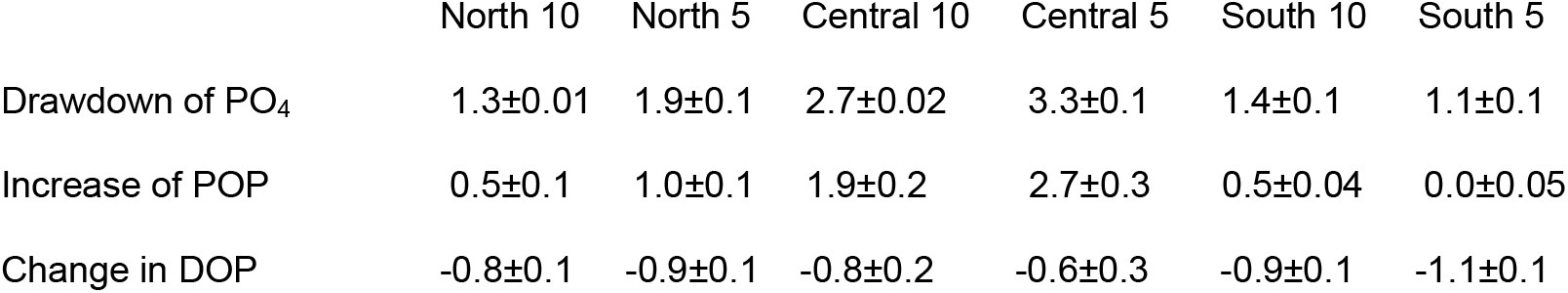
The dissolved organic phosphorus (DOP) was calculated by comparing the drawdown of PO4 (concentration at T0 minus concentration at T12) with the increase in particulate organic phosphorus (POP) (concentration at POP maximum minus concentration at T0). The calculated change in DOP is the difference between increase in POP and drawdown of PO4, either suggesting uptake or production (negative values) of DOP. All values in μmol L-1, error estimates are given as ± Standard Error.

|  | North 10 | North 5 | Central 10 | Central 5 | South 10 | South 5 |
| --- | --- | --- | --- | --- | --- | --- |
| Drawdown of PO<sub>4</sub> | 1.3±0.01 | 1.9±0.1 | 2.7±0.02 | 3.3±0.1 | 1.4±0.1 | 1.1±0.1 |
| Increase of POP | 0.5±0.1 | 1.0±0.1 | 1.9±0.2 | 2.7±0.3 | 0.5±0.04 | 0.0±0.05 |
| Change in DOP | -0.8±0.1 | -0.9±0.1 | -0.8±0.2 | -0.6±0.3 | -0.9±0.1 | -1.1±0.1 |

### Development of Chlorophyll *a* concentration

There were clear differences between the treatments in Chl *a* fluorescence with higher Chl *a* fluorescence in the ‘NP10’ compared with the ‘NP5’ treatment (two-way ANOVA, p <0.001) (Fig 3). In the community ‘Central’, there was greater than 10-fold increase in Chl *a* fluorescence in both treatments. The ‘North’ and ‘South’ communities responded very differently with a modest (less than 2-fold) increase in Chl *a* fluorescence in the ‘NP10’ treatment. In the ‘NP5’ treatment, the Chl *a* fluorescence did not increase from day 1-3 after which it decreased in the communities ‘North’ and ‘South’. At the end of the experiment, the community ‘Central’ obtained the highest Chl *a* fluorescence followed by communities ‘North’ and ‘South’ (two-way ANOVA, p <0.001). The Chl *a* concentration was determined less frequently than Chl *a* fluorescence (Fig 3), but confirmed the overall Chl *a* development, i.e. higher concentration in the ‘NP10’ compared with ‘NP5’ treatment and larger increase in Chl *a* concentration in the community ‘Central’ compared with the other communities.

**Fig 3.**
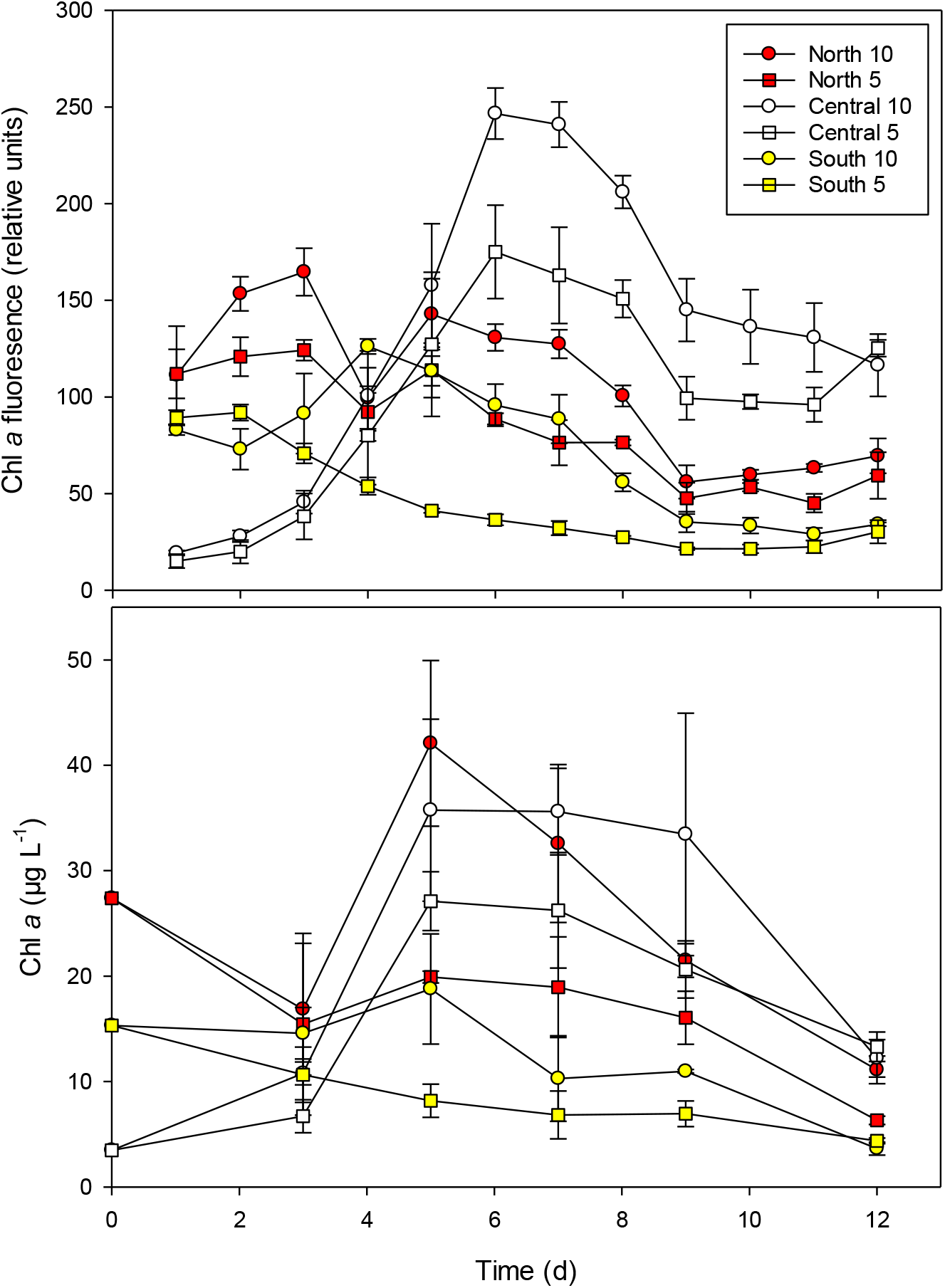
The relative Chlorophyll *a* (Chl *a*) fluorescence and Chl *a* concentration. Error bars are SE (n = 3).

### Photochemical efficiency

The photochemical efficiency (Fv/Fm) was affected by both community and NP ratio (Fig 4). Initially the Fv/Fm decreased until day 8 or 9, after which it either stabilized or increased again. Overall, it was higher in the communities ‘Central’ and ‘South’ than in the community North (Tuckey post-hoc test, p <0.001) and it was likely higher in the ‘NP5’ compared with the ‘NP10’ treatment (two-way ANOVA, p = 0.039).

**Fig 4.**
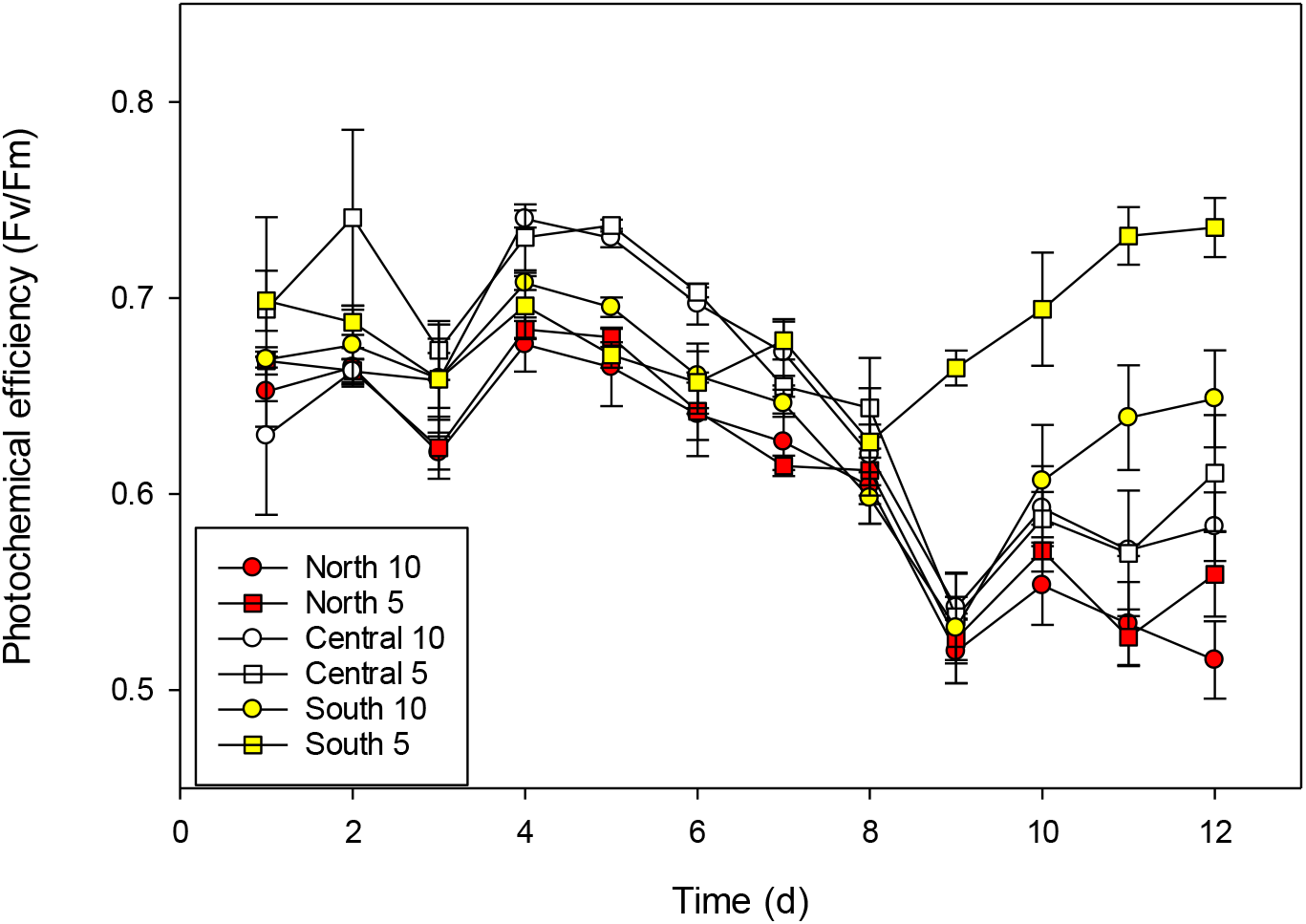
The photochemical efficiency (Fv/Fm) measured with 450 nm excitation light after 15 min dark acclimation. Error bars are SE (n = 3).

### Phytoplankton community

There were clear differences in the initial phytoplankton communities taken from the three locations that persisted throughout the experiment (Figs. 5, 6 and 7). A substantial fraction of the phytoplankton biomass was the centric diatom *Thalassiosira* sp. in the communities ‘North’ and ‘Central’. This was apparently the same species (25-35 µm Ø). The main difference between these communities was higher abundance of small (<10 µm) phytoplankton (Figs 5 and 6), mainly small flagellates and diatoms based on the microscopy counts, in the community ‘Central’ compared with the community ‘North’. This was seen as higher nanophytoplankton concentration in the community ‘Central’ compared with the other two communities (Fig. 6, Tukey post-hoc test, day 12, p ≤0.006). The development of cryptophytes was similar in all treatments (two-way ANOVA, p = 0.77), and without any effect of inorganic NP ratio (two way ANOVA, p = 0.45). In the community ‘South’, the initial biomass was dominated by dinoflagellates, in particular *Prorocentrum* sp. (Fig. 5). The share of dinoflagellates of the total biomass decreased throughout the experiment, but still made up a considerable part of the biomass at the end of the experiment (Fig 5). Picoeucaryotes were most abundant in the community ‘South’ (two-way ANOVA, p <0.001), with the highest abundance in the ‘NP5’ treatment (Tukey post-hoc test, p = 0.006), but not in the other two communities (Tukey post-hoc test, p ≥0.5). In addition to the apparently healthy phytoplankton (evaluated from the FlowCam images), an increasing share of the biomass was detritus, reaching approximately 40% of the biomass at day 12 in all treatments, except for the treatment ‘South-NP10’ where this was slightly higher at 50%.

**Fig 5.**
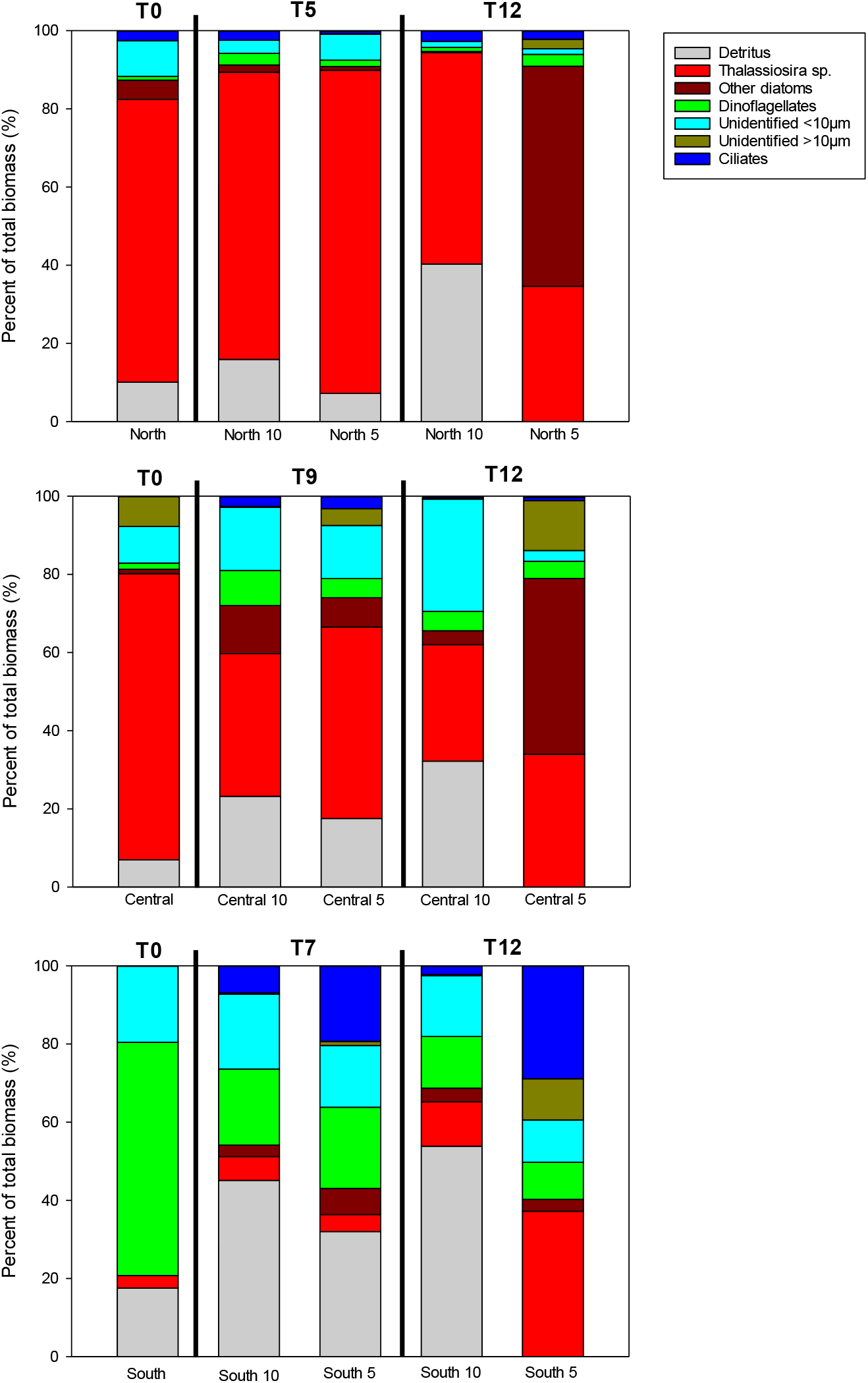
The average (n = 3) contribution of different plankton categories as percent of total biovolume. Sampling was right before the start of the experiment (T0), at the Chl *a* peak indicated by fluorescence measurements (T5, T7 and T9) and at the end of the experiment (T12).

**Fig 6.**
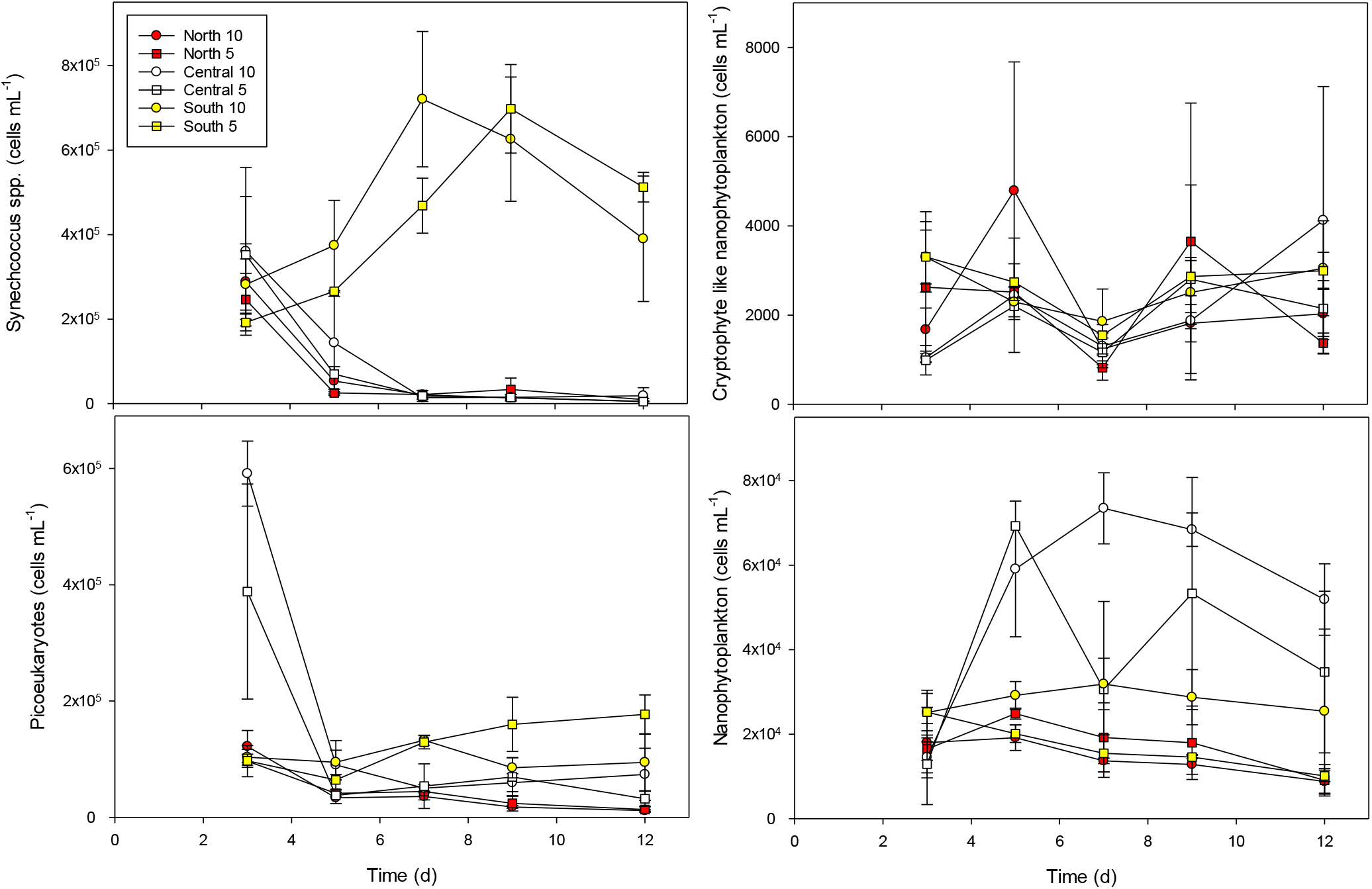
The counts on picoplankton with a flow cytometer based on auto fluorescence. Error bars are SE (n = 3).

**Fig 7.**
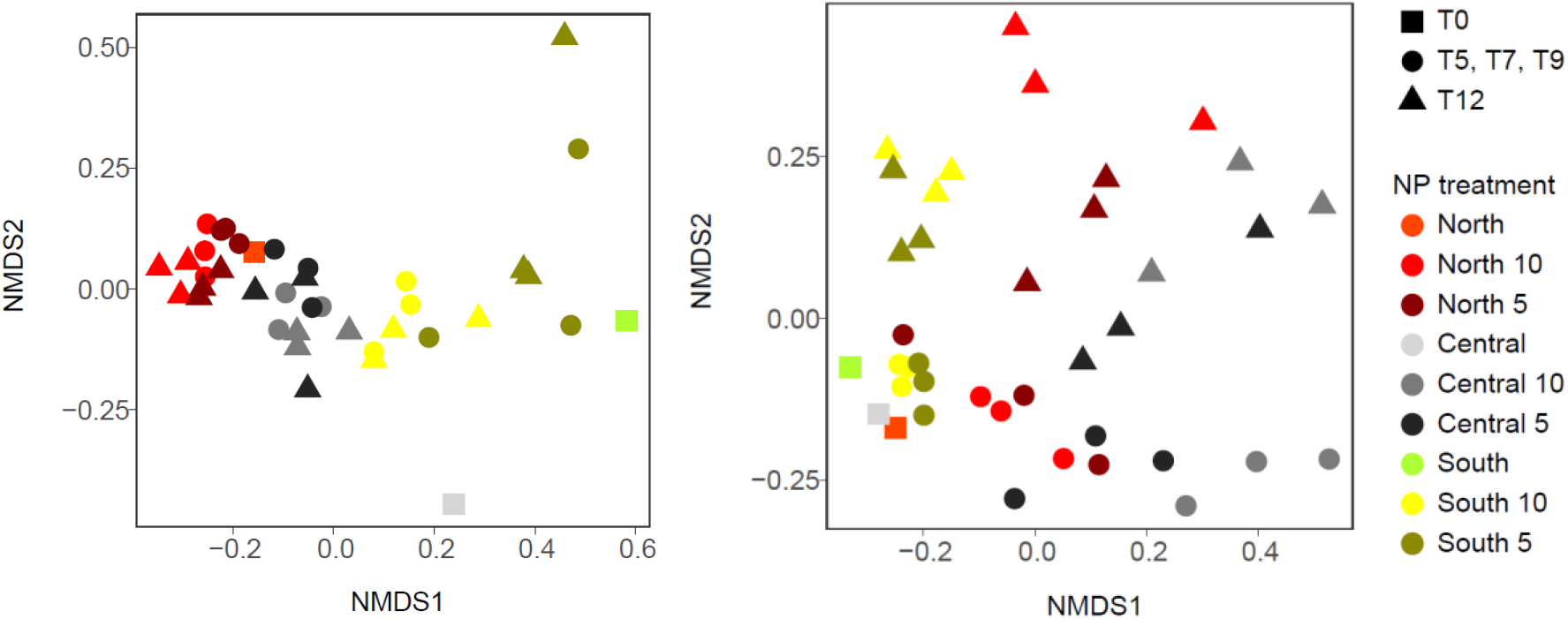
Non-metric multidimensional scaling (NMDS) plots showing the (A) phytoplankton community dynamics based on light microscopy and (B) bacterial community, based on 16S ribosomal RNA (rRNA) gene sequences dynamics, during the experiment.

The abundances of the cyanobacteria *Synechococcus* spp. were initially similar in the all three communities, but increased and leveled off at a much higher concentration in the community ‘South’, compared with the communities ‘North’ and ‘Central’ (Fig 6; Tukey post-hoc test, p <0.001). The average abundance of *Synechococcus* spp. was higher in the ‘NP5’ than in the ‘NP10’ treatment, but without a clear difference (two-way ANOVA, p = 0.09). However, based on the flow cytometer counts, the overall abundance of *Synechococcus* spp. was likely higher in the low NP treatment (ANOVA, p = 0.02) in the community ‘South’, which was similar to the relative abundance of *Synecococcus* operational taxonomic units (OTUs) (ANOVA, *p* = 0.02) at the end of the experiment.

### Heterotrophic bacteria

The initial communities of heterotrophic bacteria were similar in all three areas (Fig 7). The communities were dominated by the class Alphaproteobacteria with higher contribution of genera from the *Roseobacter* clade (genera *Amylibacter, Loktanella* and *Planktomarina*), and the class Flavobacteriia (genus *Polaribacter*), contributing 32-40 % and 20-34 % of the total relative abundance respectively (Fig. 8). Gammaproteobacteria, and Cyanobacteria were also present but with lower relative abundance (6-10 % and 4-17 %, respectively) of the OTUs. At the end of the experiment, the bacterial community composition differed between ‘North’, ‘Central’ and ‘South’ (PERMANOVA, *p* <0.001), but the overall community composition was likely not different between the NP treatments (PERMANOVA, *p* = 0.06). There were, however, differences in the community ‘South’, where the relative abundance of the genus *Candidatus Actinomarina* (Acidimicrobiia) was likely higher in the ‘NP10’ (12%) compared with the ‘NP5’ (6%) treatment (ANOVA, *p* = 0.03).

**Fig 8.**
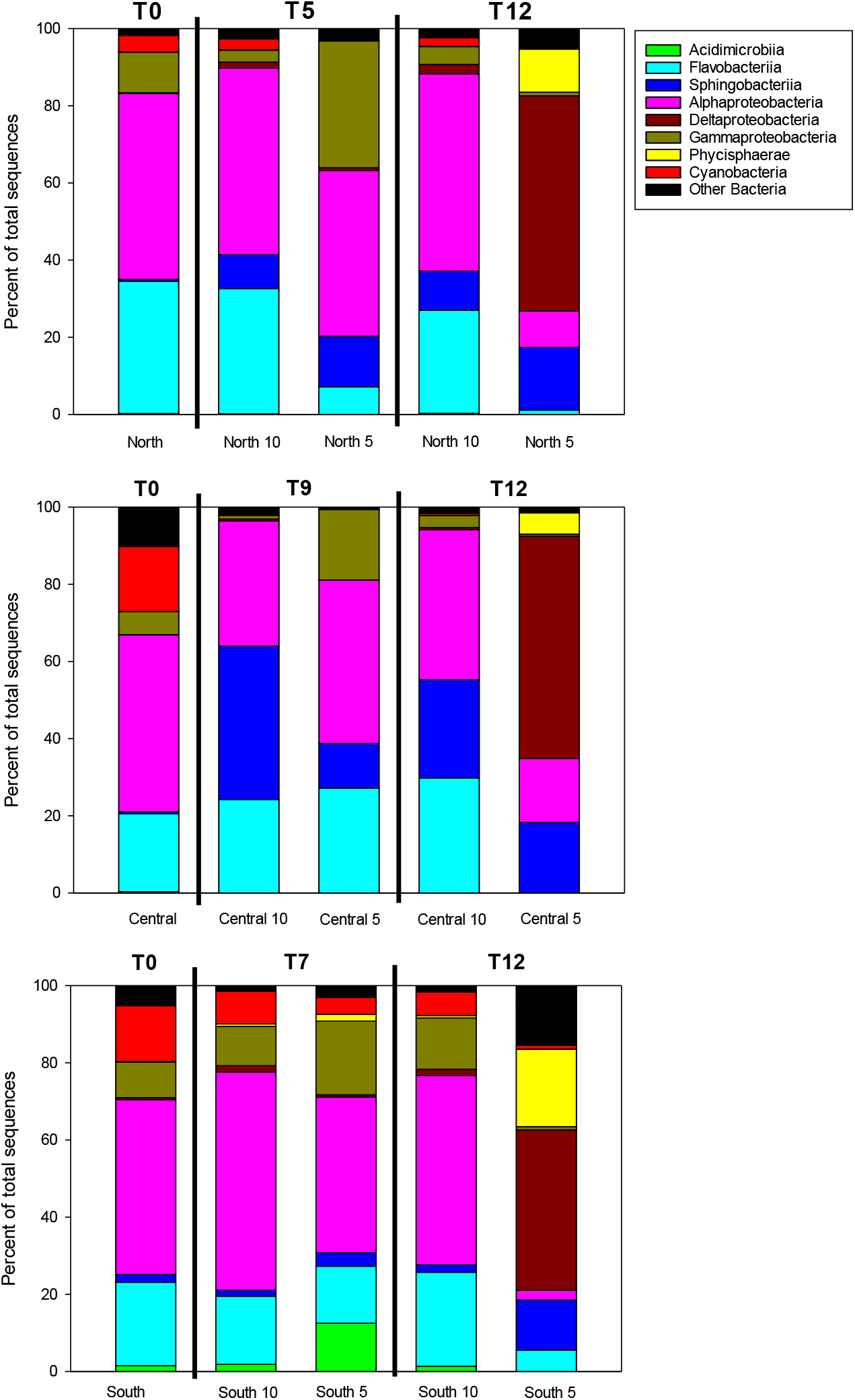
The group level Operational taxonomic units (OTUs) of bacteria. The full sequence of OTUs can be found in Supplementary Fig. S3.

### Zooplankton abundance

The abundance of zooplankton was initially different in the collected water and this difference persisted throughout the experiment (Figs. 5 and 9). Ciliates made up a large share of the biomass in the community ‘South’, whereas heterotrophic dinoflagellates (mainly *Protoperidinium* sp.; separate microscope counts not shown) were more abundant in the communities ‘North’ and ‘Central’ (Tukey post-hoc test, p ≤0.001), but no NP treatment effect was observed for microzooplankton (two-way ANOVA, p = 0.24). In addition to ciliates, more copepods (mesozooplankton) developed in the community ‘South’ compared with the communities ‘North’ and ‘Central’ (Fig. 6, Tuckey’s post-hoc test, p ≤0.005), and similar to microzooplankton, without indication of difference between the NP treatments (two-way ANOVA, p = 0.26).

**Fig 9.**
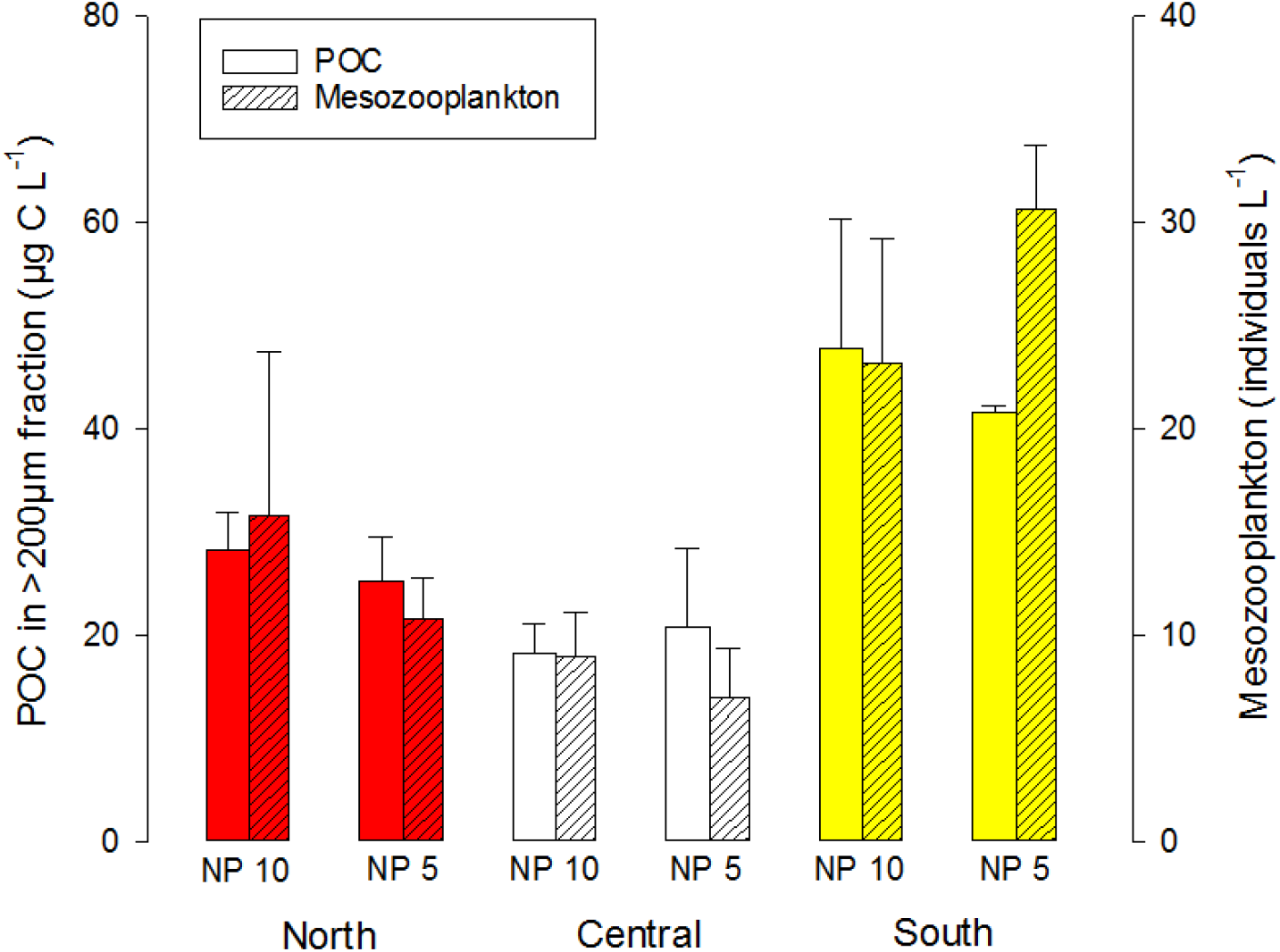
The concentration of particulate organic carbon (POC) in the >200 µm fraction and number of mesozooplankton individuals counted on the filter at the end of the experiment.

Using the initial N available (NO_3_ + PON) at the beginning of the experiment, and the PON in the >200 µm fraction at the end of the experiment, the calculated mean transfer of N to mesozooplankton was highest in the treatment ‘South-NP5’ with 2.9% followed by the treatments ‘North-NP5’ 2.6%, ‘South-NP10’ 2.4% and ‘North-NP10’ 2.2% (Table 4). In the community ‘Central’, the N transfer was considerably lower (Tukey post-hoc test, *p* <0.01) at 0.8% and 0.6% in the ‘Central-NP5’ and the ‘Central-NP10’ respectively. Overall, the N transfer efficiency to mesozooplankton was likely higher in the ‘NP5’ compared with the ‘NP10’ treatment (two-way ANOVA *p* = 0.02).

**Table 4.** The trophic transfer of nitrogen (N) into mesozooplankton during the experiment. This was calculated as the percentage of the original N pool (NO3 + PON) that was found in the mesozooplankton (>200 µm) on the last day. Error estimates are given as ± Standard Error.

| 901 |  | NP ratio 10 | NP ratio 5 |
| --- | --- | --- | --- |
| 902 | North | 2.2 % ± 0.6 | 2.6 % ± 0.4 |
| 903 | Central | 0.6 % ± 0.1 | 0.8 % ± 0.2 |
| 904 | South | 2.4 % ± 0.6 | 2.9 % ± 0.2 |

### Seston stoichiometry

The development of seston stoichiometry was clearly different depending on the plankton community composition, and there was also an effect of inorganic NP ratio on the seston NP and Chl *a*-C ratios (Fig. 10).

**Fig 10.**
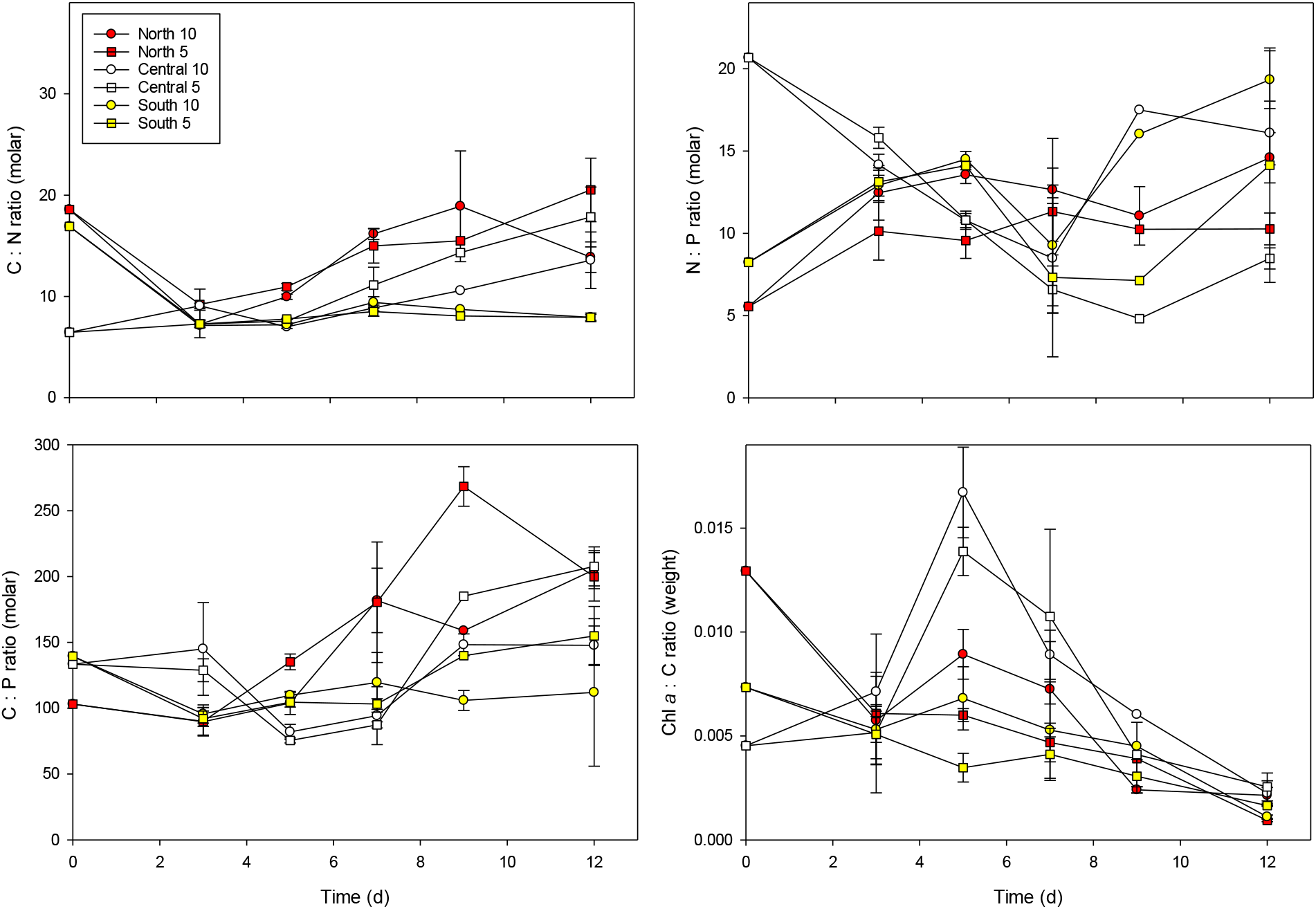
The molar stoichiometric ratios of particulate organic Carbon to Nitrogen (C : N), Carbon to Phosphorus (C : P), Nitrogen to Phosphorus (N : P) and the weight ratio of Chlorophyll *a* to Carbon (Chl *a*-C). Error bars are SE (n = 3).

The seston NP ratio increased initially in the communities ‘North’ and ‘South’ from 5 and 8 to ∼11 and 13, respectively. In the community ‘Central’ the NP ratio decreased at the start of the experiment from 21 to ∼15. The NP ratio remained stable in the community ‘North’ whereas it increased towards the end of the experiment in the communities ‘Central’ and ‘South’, with highest NP ratio in the community ‘South’ at day 12 (Fig. 10; two-way ANOVA, day 12, *p* = 0.001). After day 5, the NP ratio was consistently higher in the ‘NP10’ compared with the ‘NP5’ treatment (two-way ANOVA, day 12, *p* = 0.01). On the last day of the experiment, the average difference in seston NP ratio was 5.7 between the ‘NP10’ and the ‘NP5’ treatments.

The Chl *a*-C ratio decreased at the start of the experiment in the communities ‘North’ and ‘South’ followed by a slight increase for ‘North10’ and ‘South10’. In the community ‘Central’ there was an initial increase in the Chl *a-*C ratio to ∼0.015. For all treatments the Chl *a*-C ratio decreased after the depletion of NO_3_. There was both a community and a NP treatment effect (two way ANOVA, *p* <0.001 and *p* = 0.002, respectively), with lower Chl *a*-C ratio in the community ‘South’ compared with communities ‘North’ and ‘Central’ and higher in ‘NP10’ compared with ‘NP5’ when considering the whole experiment.

For communities ‘North’ and ‘South’, the CN ratio decreased rapidly at the onset of the experiment (from CN >15). The community ‘Central’ had an initial CN ratio close to the Redfield ratio (6.6). After the depletion of NO_3_, the CN ratio started to increase again in communities ‘North’ and ‘Central’, whereas for community ‘South’ the CN ratio remained stable. Overall, there was only a community effect on the CN ratio, with a lower ratio in community ‘South’ compared with communities ‘North’ and ‘Central’ (Tuckey post-hoc test, *p* <0.001), and no NP treatment effect (two-way ANOVA, *p* = 0.26). A similar pattern was observed for the CP ratio but with less statistical support for any difference between communities (two-way ANOVA, *p* = 0.06) and no effect of different NP ratios (two-way ANOVA, p = 0.3; Fig 10).

## Discussion

### Plankton dynamics

The phytoplankton community composition and biomass concentration were clearly different in the water that was collected from the three different locations. Inorganic N (NO_3_) was not depleted in any of the communities, suggesting that the collected phytoplankton communities were still actively growing. Based on the inorganic nutrients and Chl *a* concentrations, the phytoplankton community ‘North’ was in the most advanced growth phase (high Chl *a*, low NO_3_) followed by the community ‘South’, whereas the community ‘Central’ was in early growth phase (low Chl *a*, high NO_3_). The peak biomass was dependent on the sum of initial and added nutrients. However, phytoplankton grew exponentially at the start of the experiment in all treatments enabling comparison of the responses to different inorganic NP ratio in the three distinct communities.

Diatoms dominated the communities ‘North’ and ‘Central’ whereas dinoflagellates dominated the community ‘South’. Diatom dominance is typical for upwelling regions due to their rapid growth after upwelling events, whereas slower growing dinoflagellates typically are adapted to lower concentrations of inorganic nutrients and dominate when the water column is stratified (Reynolds 2006). Dinoflagellates have been shown to dominate in the coastal area outside the Maipo river, likely linked to the hydrological conditions here (Wieters et al. 2003). With a head start, the dinoflagellates were able to keep their initial dominant position, i.e. the faster growing diatoms were not able to out-compete them before the inorganic N pool had been depleted (Kremp et al. 2008). The two diatom dominated communities (‘North’ and ‘Central’), differed in size distribution and the relatively large (>20µm) *Thalassiosira* sp. had a more dominant role in the community ‘North’ whereas the ‘Central’ community had a larger proportion of smaller nanophytoplantkon.

The heterotrophic bacterial community composition is known to be influenced by the stage of the phytoplankton bloom and the phytoplankton community composition (Buchan et al. 2014). Despite the differences in the initial phytoplankton community, the heterotrophic bacteria communities were similar in all three locations and dominated by Alphaproteobacteria (*Roseobacter* clade) and Flavobacteriia. These groups decreased through the experiment whereas Gammaproteobacteria (Alteromonadales) increased in all the communities. Their difference in relative abundance through the experiment was probably due to the different growth phases and species composition of the phytoplankton communities. For example, the genus *Polaribacter* (Flavobacteriia) is typically abundant during active phytoplankton growth whereas Gammaproteobacteria are commonly present during the decay of phytoplankton blooms (Teeling et al. 2012). The *Roseobacter* clade, Alteromonadales and Flavobacteriia are considered the ‘master recyclers’ of algal-derived DOM during phytoplankton blooms (Chafee et al. 2017). In the community ‘South’, the proportion of Acidimicrobiia (Actinobacteria) increased in the end of the experiment. This group has been associated with cyanobacteria-derived DOM (Hugerth et al. 2015), and could have benefitted from the increase in abundance of *Synecococcus* spp in the community ‘South’.

The development of the grazer community was different between the three communities, but without any apparent effect of inorganic NP ratio. Interestingly the community with most mesozooplankton and ciliates were the dinoflagellate dominated community (‘South’). Diatoms is often considered optimal food for copepods, but the dominating dinoflagellate (*Prorocenrum* sp.) in this experiment seems also to provide a good link between primary producers and higher tropic levels through the classical food chain. For microzooplankton, there are some indications that higher P concentration of phytoplankton could benefit e.g. heterotrophic dinoflagellates (Meunier et al. 2018), but we did not find this in our short-term experiment, although there was a clear effect on seston NP ratio.

### Nutrient limitation

The primary production in the Humboldt Current is typically N-limited (Capone and Hutchins 2013; Messié and Chavez 2015), and our results clearly demonstrated N-limitation in all treatments, as the NO_3_ concentration had a direct effect on the phytoplankton biomass (iron and other trace elements had been added in excess) and the PO_4_ and DSi pools had not been depleted at the end of the experiment. There were, however, some indications of different degrees of nutrient stress in the different communities reflecting the stage of the phytoplankton community in the initial water samples.

Regenerated production with low concentrations of inorganic nutrients typically favors smaller cells due to a higher surface to volume ratio (Edwards et al. 2012), and this was observed in the community ‘South’ with higher abundance of small (0.8 – 1.5 µm) *Synechococcus* spp. and picoeucaryotes. A higher degree of regenerated production in the community ‘South’ after NO_3_ depletion was also supported by the organic (seston) CN ratio. In the communities ‘North’ and ‘Central’, the CN ratio decreased during NO_3_ uptake, followed by an increase after NO_3_ was depleted, which is expected when primary producers starts to be N-limited (Sterner and Elser 2002). In the community ‘South’, the seston CN ratio remained low (<10) throughout the experiment, suggesting that the community was less N-limited, possibly due to more effective nutrient recycling, and a similar pattern was seen for the CP ratio. That the community was less affected by N-limitation was also supported by the increase of the photochemical efficiency (Fv/Fm) in the latter half of the experiment. The Fv/Fm is affected by the phytoplankton community composition, but also by the physiological status of the cells (Suggett et al. 2009). The increase in Fv/Fm towards the end of the experiment, without any apparent changes in the community, could be an indication of less nutrient stress in the community ‘South’. The Fv/Fm was overall higher at low inorganic NP ratio, which similarly could be an indication that the communities with more rapidly depleted NO_3_ had longer time to acclimate to low nutrient conditions.

Another difference in N-availability between the communities was in the dissolved organic nitrogen (DON) pool. The apparent drawdown of DON, based on closing the N budget, was higher in the community ‘North’ than in the communities ‘Central’ and ‘South’. This drawdown was rapid and although we cannot pinpoint the bacterial group(s) responsible for the DON drawdown, several groups, such as Flavobacteriales, are known to use DON (Cottrell and Kirchman 2000). Even a single strain of bacteria is capable of depleting labile sources of nutrients quickly, even under heavy grazing pressure (Pedler et al. 2014). Given that the bacterial community was similar in the three communities at the beginning of the experiment, the higher DON utilization in the community ‘North’ could have been due to a higher proportion of labile DON produced by diatoms that was taken up by the bacterial community of this area.

### Top-down control

There were clear differences in grazing pressure between the treatments. Most notably was the more pronounced development of mesozooplankton grazing in the community ‘South’ that effectively reduced (‘NP10’ treatment) or prevented (‘NP5’ treatment) any increase in nano-and microphytoplankton. For picoplankton, however, the data suggests a different grazing regime, with higher grazing on picoplankton in the communities ‘North’ and ‘Central’. This was reflected in the rapid decrease in *Synechococcus* spp. and picoeucaryotes, which increased in the community ‘South’. In addition, the proportion of filamentous bacteria (e.g. *Lewinella*) in the diatom dominated communities was higher than in the dinoflagellate dominated community. The filamentous morphology has been suggested to be a strategy to avoid grazing (Alonso-Sáez et al. 2009; Eckert et al. 2012). We do not have direct data on bacterial grazing but the abundance of heterotrophic dinoflagellates (higher in communities ‘North’ and ‘Central’), the rapid decrease of *Synechococcus* spp. and picoeucaryotes and the relative increase of filamentous bacteria, all indicate higher grazing control of the picoplankton in the diatom dominated communities (‘North’ and ‘Central’). This could be an example of a trophic cascade, or top-down forcing (Ripple et al. 2016); if the more abundant microzooplankton were also feeding on the nano-and microzooplankton community there could be an indirect effect on the picoplankton community by reducing the grazing pressure them in the community ‘South’. This was not observed for ciliates, that were more abundant in the community ‘South’, but e.g. heterotrophic nanoflagellates are known to feed on *Synechococcus* (Dolan and Šimek 1999) and bacteria (Tsai et al. 2016).

The trophic transfer of N from phytoplankton to mesozooplankton spanned from 0.6% to 2.9% during the 12 days (Table 4). We did not measure grazing rates, and this value only represents the percent of the original N pool (inorganic and <200 µm particulate) that was incorporated into mesozooplankton (>200 µm) during the experiment, and would clearly have been higher if the mesozooplankton had not been removed before the start of the experiment. The microzooplankton biomass was not included, and assuming a biovolume to carbon conversion factor of 0.19 pg C µm^-3^ and C:N ratio of 6 for ciliates (Putt and Stoecker 1989), the highest trophic transfer from phytoplankton to grazers was 5.8%.

The Humboldt Current has one of the richest fisheries in the world, much higher than comparable Eastern Boundary Upwelling Systems (EBUS) although the primary production supporting higher trophic levels is comparable (Carr 2001; Chavez et al. 2008). The much higher fish catch off the coast of Chile and Perú is often referred to as the Humboldt paradox (Brochier et al. 2011). Most studies on this phenomenon have focused on mid-trophic level fish (i.e. wasp-waist food web) and there are data suggesting a shorter food chain supporting higher fish catches in the Humboldt Current (Chavez and Messié 2009). The results from this study point to clear differences in trophic transfer efficiency from primary producers to first level consumers (herbivores) depending on the plankton community composition with potential implications higher in the food web. The higher N transfer efficiency to mesozooplankton in the low NP treatment counteracted the lower biomass of primary producers; i.e. the lower concentration of inorganic nitrogen caused lower phytoplankton biomass, but the relative transfer to the next trophic level was higher and there was no noticeable effect on the biomass of grazers. As the initial set-up of the experiment was from a homgenious water pool, the number of microzooplankton (e.g. copepod nauplii <200 µm) can be assumed to have been the same in the two NP treatments (but different in the three communities). The results did not indicate any difference in somatic growth from micro-to mesozooplankton (<200 µm to >200µm) suggesting no difference between NP treatments in food availability for the developing mesozooplankton population. With lower phytoplankton biomass production in the low NP treatment (NP5), this provided higher trophic transfer efficiency (Fig 9). However, with a longer duration of the experiment we expect that the mesozooplankton population would have reflected the food availability.

### Stoichiometry

Our results demonstrated that part of the excess P is taken up and retained in the particular fraction resulting in a lower NP ratio in the seston, which is in consistence with other studies (Franz et al. 2012). A share of the incorporated P could be release as dissolved organic phosphorus (DOP) (Franz et al. 2012), something we did not measure directly. However, the decrease of PO_4_ and increase of POP suggested that a considerable share (average of 0.8 µmol L^-1^) of the PO_4_-P was taken up and released as DOP, but this share was not affected by the different NP ratios. For the phytoplankton community composition there was an effect. In all treatments there was excess P left at the end of the experiment, but the two diatom-dominated communities (‘North’ and ‘Central’) demonstrated higher capacity of P uptake. In these two communities, the P uptake stopped in the ‘NP10’ treatment (with more added NO_3_) after NO_3_ had been depleted whereas in the ‘NP5’ treatment it continued after NO_3_ depletion, reducing the initial difference in the inorganic P pool with 70-80%. In the community (‘South’), where dinoflagellates and *Synechococcus* spp. were more predominant, the initial difference in PO_4_ was still present at the end of the experiment. A possible explanation is that diatoms are better at drawing down and storing excess P. The communities with more diatoms could also have produced and excreted more labile carbon, which could have enhanced the bacterial P uptake (Stets and Cotner 2008). Storage capacity is affected by the overall community size structure (surface to volume ratio). High surface ratio is better for capturing scarce nutrients; whereas low surface to volume ratio increase the intracellular storage space. The higher proportion of smaller cells in the ‘South’ community could partly explain the lower capacity to take up excess P. However, it does not explain the apparent difference in P uptake capabilities after NO_3_ exhaustion between the two NP treatments and this remains an open question.

Dissolved silicate (DSi) is mainly taken up by diatoms and some smaller groups such as silicoflagellates, thus potentially affecting the phytoplankton community composition (Egge and Aksnes 1992). The DSi is incorporated into biogenic silicate in the diatom cell wall, i.e. frustules, and therefore DSi limitation favors non-siliceous phytoplankton. All treatments had excess DSi and it is therefore unlikely to have affected the phytoplankton community composition. As with PO_4_, the drawdown of DSi was highest in the diatom dominated community ‘Central’, which had highest Chl *a* fluorescence in the end of the experiment, which is not surprising as neither dinoflagellates nor *Synechococcus* spp. take up DSi.

### Decreasing inorganic NP ratio

One of the central questions with a decreasing inorganic NP ratio in the upwelling water is: what happens with the increasing oversupply of P? As the upwelling water is transported off-shore, will the elevated PO_4_ concentration increase the proportion of diazotrophs that can fix N from the atmosphere and will there be changes to the bacterial community with an increasing pool of bioavailable P? The present study was too short to answer these questions, but it gives a clear indication that the plankton community composition will affect the ability to take up excess nutrients such as PO_4_ and DSi. There are some studies of the effect of decreasing NP ratio on diazotrophs, most demonstrating no effect for N fixation, at least in the short term (Wasmund et al. 2015). In our experiment, the cyanobacteria *Synechococcus* spp. seem to have benefitted from lower NP ratio when the grazing pressure was low; *Synechococcus* is not considered to be as efficient at N-fixation as e.g. *Trichodesmium* sp, but some strains may have this ability (Fay 1992). Other studies have demonstrated that decreasing NP ratio benefits phytoplankton taxa that are of reduced food quality for mesozooplankton, with potential consequences for trophic transfer (Hauss et al. 2012). We observed slightly higher trophic transfer efficiency at low NP ratio, but with lower overall biomass, and it is clear that changes to the inorganic NP ratio may have consequences for the primary producers as well as for higher trophic levels in the food web. With an almost 1:1 shift in the seston NP stoichiometry, any decrease in the inorganic NP ratio will have repercussions for the diet of the grazing community (Sterner and Elser 2002).

In conclusion, the present experiment strengthened the hypothesis that a reduction in the inorganic NP ratio will directly affect seston stoichiometry. We did also find some indication of slight shift in the community, giving some support for the hypothesis that a shift in the inorganic NP ratio will affect the plankton community structure. This might have been more pronounced if our experiment has lasted longer and including pulses of nutrient additions simulating upwelling. Our main finding was, however, the importance of the plankton community composition, supporting our hypothesis of higher variability between communities than between NP treatments. Any long term change in plankton community structure will likely have greater impact on ecosystem functioning than any direct effects of a decreasing inorganic NP ratio in the Humboldt Current ecosystem.

## Acknowledgements

Financial support for this study came from the EU FP7 research infrastructure initiative ASSEMBLE (grant agreement no. 227799). Further support came from the Academy of Finland (decision no 259164) and the Walter and Andrée de Nottbeck foundation (KS and TC).

We would like to thank Sylvain Faugeron, the local ASSEMBLE coordinator, and the staff at Las Cruses Marine Biological Station, in particular Randy Finke who helped us with the collection of the water. We want to thank Paola Reinoso for performing the nutrient analysis, and express our sincere gratitude towards the Viu family for their hospitality.

## Author contributions

KS and VM devised the experiment, which was carried out by MTCG, TL, AM, FD and KS. EER and MTCG did the bacterial community analysis, PvD, TL and KS the phytoplankton community analysis and MTCG the community statistics. NS was responsible for the inorganic nutrient measurements. KS wrote the manuscript with input from all the co-authors.

## Additional Information

The authors declare no competing interests.

## Data availability

All data generated or analyzed during this study are included in this published article (and its supplementary information files).

## References

Aldunate M, De la Iglesia R, Bertagnolli AD, Ulloa O (2018) Oxygen modulates bacterial community composition in the coastal upwelling waters off central Chile. Deep Sea Research Part II: Topical Studies in Oceanography doi10.1016/j.dsr2.2018.02.001

Alonso-Sáez L, Unanue M, Latatu A, Azua I, Ayo B, Artolozaga I, Iriberri J (2009) Changes in marine prokaryotic community induced by varying types of dissolved organic matter and subsequent grazing pressure. J Plankton Res 31: 1373–1383

Anabalón V, Morales C, Escribano R, Varas MA (2007) The contribution of nano-and micro-planktonic assemblages in the surface layer (0–30 m) under different hydrographic conditions in the upwelling area off Concepción, central Chile. Progress in Oceanography 75: 396–414

Anabalón V, Morales C, González H, Menschel E, Schneider W, Hormazabal S, Valencia L, Escribano R (2016) Micro-phytoplankton community structure in the coastal upwelling zone off Concepción (central Chile): Annual and inter-annual fluctuations in a highly dynamic environment. Progress in Oceanography 149: 174–188

Anderson M, Gorley RN, Clarke RK (2008) Permanova+ for Primer: Guide to Software and Statisticl Methods. Primer-E Limited

Anderson MJ (2001) A new method for non-parametric multivariate analysis of variance. Austral ecology 26: 32–46

Anderson MJ (2006) Distance-based tests for homogeneity of multivariate dispersions. Biometrics 62: 245–253

Anderson MJ, Ellingsen KE, McArdle BH (2006) Multivariate dispersion as a measure of beta diversity. Ecol Lett 9: 683–693

Atlas EL, Gordon L, Hager S, Park PK (1971) A practical manual for use of the Technicon Autoanalyzer in seawater nutrient analyses. Oregon State University, Department of Oceanography

Brochier T, Lett C, Fréon P (2011) Investigating the ‘northern Humboldt paradox’from model comparisons of small pelagic fish reproductive strategies in eastern boundary upwelling ecosystems. Fish and Fisheries 12: 94–109

Buchan A, LeCleir GR, Gulvik CA, González JM (2014) Master recyclers: features and functions of bacteria associated with phytoplankton blooms. Nature Reviews Microbiology 12: 686

Capone DG, Hutchins DA (2013) Microbial biogeochemistry of coastal upwelling regimes in a changing ocean. Nature Geosci 6: 711–717

Carr M-E (2001) Estimation of potential productivity in Eastern Boundary Currents using remote sensing. Deep Sea Research Part II: Topical Studies in Oceanography 49: 59–80

Chafee M, Fernàndez-Guerra A, Buttigieg PL, Gerdts G, Eren AM, Teeling H, Amann RI (2017) Recurrent patterns of microdiversity in a temperate coastal marine environment. The ISME journal 12: 237

Chavez FP, Bertrand A, Guevara-Carrasco R, Soler P, Csirke J (2008) The northern Humboldt Current System: Brief history, present status and a view towards the future. Progress in Oceanography 79: 95–105

Chavez FP, Messié M (2009) A comparison of eastern boundary upwelling ecosystems. Progress in Oceanography 83: 80–96

Collado-Fabbri S, Vaulot D, Ulloa O (2011) Structure and seasonal dynamics of the eukaryotic picophytoplankton community in a wind-driven coastal upwelling ecosystem. Limnol Oceanogr 56: 2334–2346

Cottrell MT, Kirchman DL (2000) Natural assemblages of marine proteobacteria and members of the Cytophaga-Flavobacter cluster consuming low-and high-molecular-weight dissolved organic matter. Appl Env Microbiol 66: 1692–1697

Dolan JR, Šimek K (1999) Diel periodicity in Synechococcus populations and grazing by heterotrophic nanoflagellates: analysis of food vacuole contents. Limnol Oceanogr 44: 1565–1570

Eckert EM, Salcher MM, Posch T, Eugster B, Pernthaler J (2012) Rapid successions affect microbial N-acetyl-glucosamine uptake patterns during a lacustrine spring phytoplankton bloom. Env Microbiol 14: 794–806

Edgar RC (2013) UPARSE: highly accurate OTU sequences from microbial amplicon reads. Nature methods 10: 996

Edgar RC, Haas BJ, Clemente JC, Quince C, Knight R (2011) UCHIME improves sensitivity and speed of chimera detection. Bioinformatics 27: 2194–2200

Edwards KF, Thomas MK, Klausmeier CA, Litchman E (2012) Allometric scaling and taxonomic variation in nutrient utilization traits and maximum growth rate of phytoplankton. Limnol Oceanogr 57: 554–566

Egge JK, Aksnes DL (1992) Silicate as regulating nutrient in phytoplankton competition. Mar Ecol Prog Ser 83: 281–289

Eggers SL, Lewandowska AM, Barcelos e Ramos J, Blanco-Ameijeiras S, Gallo F, Matthiessen B (2014) Community composition has greater impact on the functioning of marine phytoplankton communities than ocean acidification. Global Change Biol 20: 713–723

Einsele W (1938) Über chemische und kolloidchemische Vorgänge in Eisen-Phosphat-Systemen unter limnochemischen und limnogeologischen Gesichtspunkten. Arch Hydrobiol 33: 361–387

Fay P (1992) Oxygen relations of nitrogen fixation in cyanobacteria. Microbiological reviews 56: 340–373

Fernandez C, Farías L, Ulloa O (2011) Nitrogen fixation in denitrified marine waters. PloS One 6: e20539

Franz J, Hauss H, Sommer U, Dittmar T, Riebesell U (2012) Production, partitioning and stoichiometry of organic matter under variable nutrient supply during mesocosm experiments in the tropical Pacific and Atlantic Ocean. Biogeosciences 9: 4629–4643

Guillard RRL (1975) Culture of phytoplankton for feeding marine invertebrates. In: Smith WL, Chanley MH (eds) Culture of marine invertebrate animals. Plenum Press, New York, pp 26–60

Hamersley MR, Turk K, Leinweber A, Gruber N, Zehr J, Gunderson T, Capone D (2011) Nitrogen fixation within the water column associated with two hypoxic basins in the Southern California Bight. Aquat Microb Ecol 63: 193–205

Hauss H, Franz JM, Sommer U (2012) Changes in N: P stoichiometry influence taxonomic composition and nutritional quality of phytoplankton in the Peruvian upwelling. J Sea Res 73: 74–85

Hugerth LW, Larsson J, Alneberg J, Lindh MV, Legrand C, Pinhassi J, Andersson AF (2015) Metagenome-assembled genomes uncover a global brackish microbiome. Genome biology 16: 279

Kalvelage T, Jensen MM, Contreras S, Revsbech NP, Lam P, Günter M, LaRoche J, Lavik G, Kuypers MM (2011) Oxygen sensitivity of anammox and coupled N-cycle processes in oxygen minimum zones. PLoS One 6: e29299

Kalvelage T, Lavik G, Lam P, Contreras S, Arteaga L, Löscher CR, Oschlies A, Paulmier A, Stramma L, Kuypers MM (2013) Nitrogen cycling driven by organic matter export in the South Pacific oxygen minimum zone. Nature Geosci 6: 228–234

Keeling RF, Körtzinger A, Gruber N (2010) Ocean deoxygenation in a warming world. Ann Rev Mar Sci 2: 199–229

Kirchman DL (2012) Processes in microbial ecology. Oxford University Press

Kremp A, Tamminen T, Spilling K (2008) Dinoflagellate bloom formation in natural assemblages with diatoms: nutrient competition and growth strategies in Baltic spring phytoplankton. Aquat Microb Ecol 50: 181–196

Kuypers MM, Lavik G, Woebken D, Schmid M, Fuchs BM, Amann R, Jørgensen BB, Jetten MS (2005) Massive nitrogen loss from the Benguela upwelling system through anaerobic ammonium oxidation. Proc Natl Acad Sci 102: 6478–6483

Landolfi A, Dietze H, Koeve W, Oschlies A (2013) Overlooked runaway feedback in the marine nitrogen cycle: the vicious cycle. Biogeosciences 10: 1351–1363

Manly BF (2006) Randomization, bootstrap and Monte Carlo methods in biology. CRC press

Martin M (2011) Cutadapt removes adapter sequences from high-throughput sequencing reads. EMBnet journal 17: pp. 10–12

Messié M, Chavez FP (2015) Seasonal regulation of primary production in eastern boundary upwelling systems. Progress in Oceanography 134: 1–18

Meunier CL, Alvarez-Fernandez S, Cunha-Dupont AÖ, Geisen C, Malzahn AM, Boersma M, Wiltshire KH (2018) The craving for phosphorus in heterotrophic dinoflagellates and its potential implications for biogeochemical cycles. Limnol Oceanogr 63: 1774–1784

Montecino V, Lange CB (2009) The Humboldt Current System: Ecosystem components and processes, fisheries, and sediment studies. Progress in Oceanography 83: 65–79

Narváez DA, Poulin E, Leiva G, Hernández E, Castilla JC, Navarrete SA (2004) Seasonal and spatial variation of nearshore hydrographic conditions in central Chile. Cont Shelf Res 24: 279–292

Ochoa N, Taylor MH, Purca S, Ramos E (2010) Intra-and interannual variability of nearshore phytoplankton biovolume and community changes in the northern Humboldt Current system. J Plankton Res 32: 843–855

Pedler BE, Aluwihare LI, Azam F (2014) Single bacterial strain capable of significant contribution to carbon cycling in the surface ocean. Proc Natl Acad Sci 111: 7202–7207

Putt M, Stoecker DK (1989) An experimentally determined carbon : volume ratio for marine “olilgotrichous” ciliates from estuarine and coastal waters. Limnol Oceanogr 34: 1097–1103

Quast C, Pruesse E, Yilmaz P, Gerken J, Schweer T, Yarza P, Peplies J, Glöckner FO (2012) The SILVA ribosomal RNA gene database project: improved data processing and web-based tools. Nucleic acids research 41: D590–D596

Reynolds CS (2006) Ecology of phytoplankton. Cambridge University Press, Cambridge

Ripple WJ, Estes JA, Schmitz OJ, Constant V, Kaylor MJ, Lenz A, Motley JL, Self KE, Taylor DS, Wolf C (2016) What is a trophic cascade? Trends Ecol Evol 31: 842–849

Schloss PD, Westcott SL, Ryabin T, Hall JR, Hartmann M, Hollister EB, Lesniewski RA, Oakley BB, Parks DH, Robinson CJ (2009) Introducing mothur: open-source, platform-independent, community-supported software for describing and comparing microbial communities. Appl Env Microbiol 75: 7537–7541

Schmidtko S, Stramma L, Visbeck M (2017) Decline in global oceanic oxygen content during the past five decades. Nature 542: 335

Solórzano L, Sharp JH (1980) Determination of total dissolved phosphorus and particulate phosphorus in natural waters. Limnol Oceanogr 25: 754–758

Sterner RW, Elser JJ (2002) Ecological Stoichiometry. Princeton University Press, Princeton

Stets EG, Cotner JB (2008) The influence of dissolved organic carbon on bacterial phosphorus uptake and bacteria-phytoplankton dynamics in two Minnesota lakes. Limnol Oceanogr 53: 137–147

Stramma L, Schmidtko S, Levin LA, Johnson GC (2010) Ocean oxygen minima expansions and their biological impacts. Deep Sea Res Pt I 57: 587–595

Strub PT, James C, Montecino V, Rutllant JA, Blanco JL (2019) Ocean circulation along the southern Chile transition region (38°–46° S): Mean, seasonal and interannual variability, with a focus on 2014–2016. Progress in Oceanography in press

Suggett DJ, Moore CM, Hickman AE, Geider RJ (2009) Interpretation of fast repetition rate (FRR) fluorescence: signatures of phytoplankton community structure versus physiological state. Mar Ecol Prog Ser 376: 1–19

Teeling H, Fuchs BM, Becher D, Klockow C, Gardebrecht A, Bennke CM, Kassabgy M, Huang S, Mann AJ, Waldmann J (2012) Substrate-controlled succession of marine bacterioplankton populations induced by a phytoplankton bloom. Science 336: 608–611

Tsai A-Y, Gong G-C, Chao CF (2016) Contribution of Viral Lysis and Nanoflagellate Grazing to Bacterial Mortality at Surface Waters and Deeper Depths in the Coastal Ecosystem of Subtropical Western Pacific. Estuaries and coasts 39: 1357–1366

Vargas CA, Martínez RA, Cuevas LA, Pavez MA, Cartes C, González HE, Escribano R, Daneri G (2007) The relative importance of microbial and classical food webs in a highly productive coastal upwelling area. Limnol Oceanogr 52: 1495–1510

Wasmund N, Struck U, Hansen A, Flohr A, Nausch G, Grüttmüller A, Voss M (2015) Missing nitrogen fixation in the Benguela region. Deep Sea Res Pt I 106: 30–41

Wieters EA, Kaplan D, Navarrete S, Sotomayor A, Largier J, Nielsen KJ, Veliz F (2003) Alongshore and temporal variability in chlorophyll a concentration in Chilean nearshore waters. Mar Ecol Prog Ser 249: 93–105

Yang S, Gruber N, Long MC, Vogt M (2017) ENSO-Driven Variability of Denitrification and Suboxia in the Eastern Tropical Pacific Ocean. Global Biogeochem Cy 31: 1470–1487

Zhang J, Kobert K, Flouri T, Stamatakis A (2013) PEAR: a fast and accurate Illumina Paired-End reAd mergeR. Bioinformatics 30: 614–620

